# Identification of Novel Scaffold Proteins for Improved Endogenous Engineering of Extracellular Vesicles

**DOI:** 10.1101/2023.05.17.541095

**Authors:** Wenyi Zheng, Julia Rädler, Helena Sork, Zheyu Niu, Samantha Roudi, Jeremy Bost, André Görgens, Ying Zhao, Doste Mamand, Xiuming Liang, Oscar Wiklander, Taavi Lehto, Dhanu Gupta, Joel Z. Nordin, Samir EL Andaloussi

**Affiliations:** Division of Biomolecular and Cellular Medicine, Department of Laboratory Medicine, Karolinska Institutet, Huddinge, Sweden; Institute of Technology, University of Tartu, Tartu, Estonia; Institute for Transfusion Medicine, University Hospital Essen, University of Duisburg-Essen, Essen, Germany; EVOX Therapeutics Limited, Oxford, United Kingdom; Clinical Research Center, Karolinska University Hospital, Stockholm, Sweden

**Author notes:** Equal contribution (Wenyi Zheng, Julia Rädler).

**Keywords:** exosome, extracellular vesicles, tetraspanin, endogenous loading

## Abstract

Extracellular vesicles (EVs) are gaining ground as next-generation drug delivery modalities. Genetic fusion of the protein of interest to a scaffold protein with high EV-sorting ability represents a robust cargo loading strategy. To address the paucity of such scaffold proteins we conducted a large-scale comparative study involving 244 candidate proteins. Their EV-sorting potential was evaluated using a simple but reliable assay that can distinguish intravesicular cargo proteins from surface and non-vesicular proteins. Notably, 24 proteins with conserved EV-sorting abilities across five types of producer cells were identified. Most of these are first to be reported including TSPAN2 and TSPAN3, which emerged as lead candidates, outperforming the well-known CD63 scaffold. Importantly, these engineered EVs show promise as delivery vehicles as demonstrated by *in vitro* and *in vivo* internalization studies with luminal cargo proteins as well as surface display of functional domains. The discovery of these novel scaffolds provides a new platform for EV-based engineering.

## INTRODUCTION

Extracellular vesicles (EVs) are membrane-enclosed particles that are secreted by most types of cells^1–3^. By harboring diverse macromolecules, EVs are important mediators of intercellular communication owing to their intrinsic tropism and protection of their luminal contents from rapid degradation. In combination with favorable safety profiles, EVs have gained tremendous attention as a potential next-generation therapeutic modality for a wide range of diseases^4, 5^.

Harnessing EVs for therapeutic applications relies first and foremost on their contents. In the simplest scenario, EVs are inherently packed with therapeutic molecules from their source cell^6–9^. For instance, mesenchymal stem cell-derived EVs have repeatedly been shown to reflect the regenerative and immunomodulatory properties of their parental cells^10^. EVs can also be deliberately loaded with molecules of interest through exogenous or endogenous means. Exogenous loading is performed on pre-isolated EVs using physical methods such as sonication, electroporation or chemical conjugation. This approach is largely restricted to small payloads including miRNAs and low molecular weight chemicals and is associated with technical challenges related to RNA precipitation and physical impairment or aggregation of EVs^11, 12^. Larger payloads (e.g., proteins) are often loaded endogenously in the course of EV biogenesis in producing cells^13–15^. Typically, producer cells are genetically instructed to overexpress the protein of interest fused to an EV-sorting protein, thereby boosting endogenous sorting of the cargo protein. This strategy can direct molecules to the surface or the lumen of EVs^16^. In contrast to surface display, luminal loading prevents premature dissociation/degradation of the cargo and is therefore the approach of choice for molecules that are prone to degradation and operate in the cytosol or nucleus of recipient cells. We and others have demonstrated the applicability of endogenous loading by incorporating protein therapeutics like super-repressor IκB and receptor protein decoys into or onto EVs^17–22^. Additionally, this approach allows for indirect loading of RNA therapeutics by fusion of RNA-binding proteins to EV-sorting proteins^23–26^.

Although a versatile strategy, endogenous loading is essentially determined by the abundance of the sorting protein in an EV population. This refers not only to the levels it can reach per EV, but more importantly to its presence across different EV subpopulations. Generally, EV populations are heterogenous pools that to date the field struggles to characterize and physically separate into distinct subpopulations^27, 28^, which complicates endogenous loading strategies. On the upside, 213 proteins were found to be conserved across EVs from 60 different cell types from the National Cancer Institute (NCI-60), therefore lending themselves as potential EV-sorting candidates^29^. However, until now only a few proteins have been well-characterized for loading proteins into EVs. Most of these are multi-pass transmembrane proteins belonging to the tetraspanin superfamily, such as CD9, CD63 and CD81^30^. In reflection of the heterogeneity of EVs, over-expression of CD63-GFP fusion protein resulted in 51% GFP-positive particles at most^31–33^. Efforts have been made to identify novel EV-sorting candidates showing promise for PTGFRN and BASP1^32^ as well as TSPAN14^33^. These studies focused on low-throughput, GFP-centered quantification methods and included a maximum of 14 candidate proteins.

To obtain a more comprehensive picture and in the hopes of identifying novel EV-sorting proteins, we conducted a semi high-throughput comparative study including 244 potential candidates. Throughout the entire study, the tetraspanins TSPAN2, TSPAN3 and CD63 consistently emerged as highly efficient EV-sorting proteins showing robust luminal loading profiles across different producer cell types. Furthermore, TSPAN2- and TSPAN3-engineered EVs showed great potential as delivery modalities not only ascertained by efficient uptake *in vitro* and *in vivo* but also owing to ample engineering possibilities (i.e., surface display and luminal cargo loading). Therefore, we believe that this discovery provides a steppingstone for endogenous engineering approaches to load cargo into or onto EVs, thereby enabling potential therapeutic applications.

## RESULTS

### A simple assay for screening EV-sorting proteins

In search of novel and efficient EV-sorting proteins, a list of candidates was compiled on basis of literature review and proteomics databases. Potential candidates were derived from either (1) proteins found to be enriched in EVs across the NCI-60 cells^29^, (2) proteins abundant in EVs produced by HEK-293T cells^34^, (3) reported EV-sorting proteins as references to previous studies^32, 33^, and (4) all proteins in the tetraspanin superfamily. Proteins larger than 130 kDa were excluded to facilitate overexpression/engineering. A total of 244 candidates with a median size of 38 kDa were included, of which 129 were non- and 115 were transmembrane proteins (Figure 1a; Table S1).

**Figure 1.**
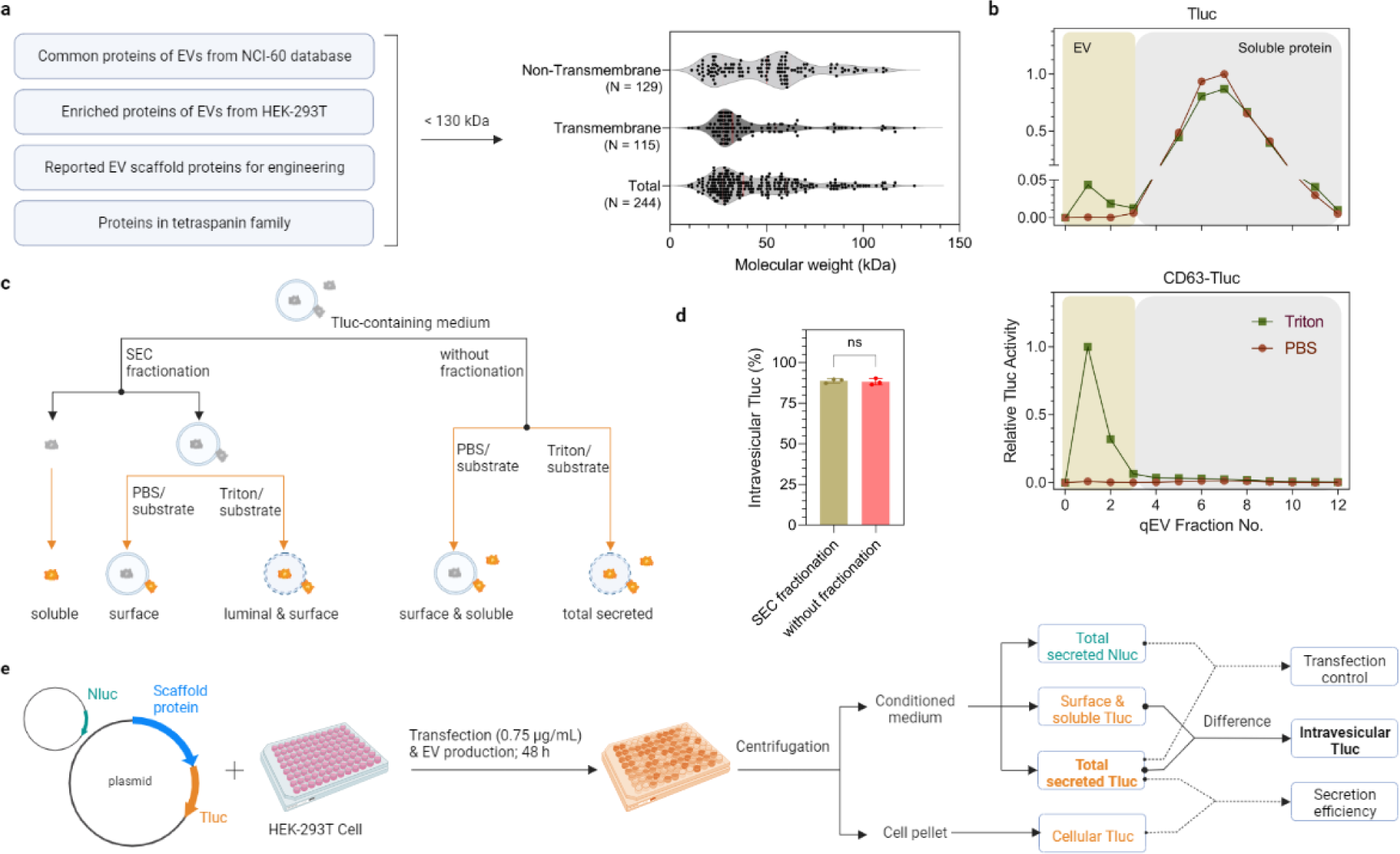
A bioluminescence screening protocol for quantification of luminal cargo proteins in EVs. (a) Selection criteria and overview of EV-sorting protein candidates. (b) Tluc-based SEC elution profiles of conditioned media from cells expressing Tluc or CD63-Tluc. Tluc activity in each fraction was quantified directly (group PBS) or after membrane lysis (group Triton) and normalized to the fraction with the highest signal. EVs and soluble proteins were recovered in fractions 0-3 and 4-12, respectively. (c) Scheme of differentiating Tluc forms in conditioned media. (d) Percentage of intravesicular Tluc for CD63-Tluc using fractionated and unfractionated media. Data were analyzed using student T test and are shown as mean ± standard deviation of three biological replicates. ns: not significant. (e) Outline of the screening procedure and data analyses. HEK-293T cells were grown in 96-well microplates and co-transfected with Tluc fusion plasmid and Nluc plasmid. Cell cultures were centrifuged and Tluc activity was measured in the cell pellet and conditioned media. Nluc activity was only quantified in the conditioned media.

To assess the luminal loading ability of the candidates into EVs, we developed an assay based on the luciferase reporter ThermoLuc (Tluc; 60.5 kDa)^18^. In brief, Tluc was fused to the C termini of all candidates, bearing in mind that N termini are usually the site for signal peptides and post-translational modifications. The plasmids encoding the fusion proteins were transfected into human embryonic kidney epithelial (HEK-293T) cells. After 48 hours, the conditioned media were collected and further processed prior to bioluminescence measurements. Initially, to evaluate the feasibility of this assay, the conditioned media of cells expressing Tluc alone and CD63-Tluc were analyzed. Both were fractionated with size exclusion chromatography (SEC) columns to separate vesicles from free proteins (Figure 1b)^35^. The fractions were treated either with PBS to determine soluble/surface-associated Tluc or the detergent Triton X-100 to detect total secreted Tluc (Figure 1c). Compared to Tluc alone, fusion with CD63 resulted in a prominent shift of Tluc towards the EV fraction (Figure 1b). Notably, Tluc activity in the EV fractions was only detected upon membrane lysis, indicating that Tluc substrate is unable to cross the EV membrane and react with luminal luciferase. This implies that SEC fractionation is dispensable for quantifying luminal proteins. This is further supported by comparison of unfractionated and fractionated media of CD63-Tluc-expressing cells, which revealed no significant differences in the percentage of intravesicular Tluc (Figure 1d). Taken together, these data show that this assay can be used in a high-throughput format to identify potential EV-loading scaffolds.

Moving forward, the principle for screening all 244 candidates was the same as above. For downstream analyses, the proteins were primarily evaluated on the absolute amount of intravesicular Tluc, derived from the difference in Tluc signal detected with and without membrane lysis, or the relative percentage thereof (Figure 1e). Information on fusion protein expression was obtained by measuring Tluc in the EV-producing cells. Additionally, when specified, the data was normalized to a transfection control in the form of a plasmid encoding NanoLuc (Nluc) luciferase that was spiked into the transfection mixture to account for possible transfection variations.

### Screening identifies 31 novel EV-sorting proteins

Screening the 244 candidates in HEK-293T cells revealed no obvious correlation between intravesicular Tluc and cellular or total secreted Tluc (Figure S1a, Figure S1b), indicating that neither cellular expression nor overall secretion fully predicts EV-sorting ability. For most candidates the percentage of intravesicular Tluc was below zero, which is unexpected but might be attributable to attenuated enzyme activity and/or photon lifetime in the presence of the detergent Triton. Nevertheless, it provided a reasonable and practical cut-off for proteins with EV-sorting ability. According to this definition, a total of 36 proteins were found to exhibit EV-sorting ability in HEK-293T cells (Figure 2a). Among these were five known EV-sorting proteins including three canonical EV markers (CD9, CD63, CD81), one recently identified protein (PTGFRN)^32^, and the viral glycoprotein gag, thereby substantiating the validity of our screening protocol. To the best of our knowledge, the remaining 31 proteins capable of luminal cargo loading into EVs are first to be reported here.

**Figure 2.**
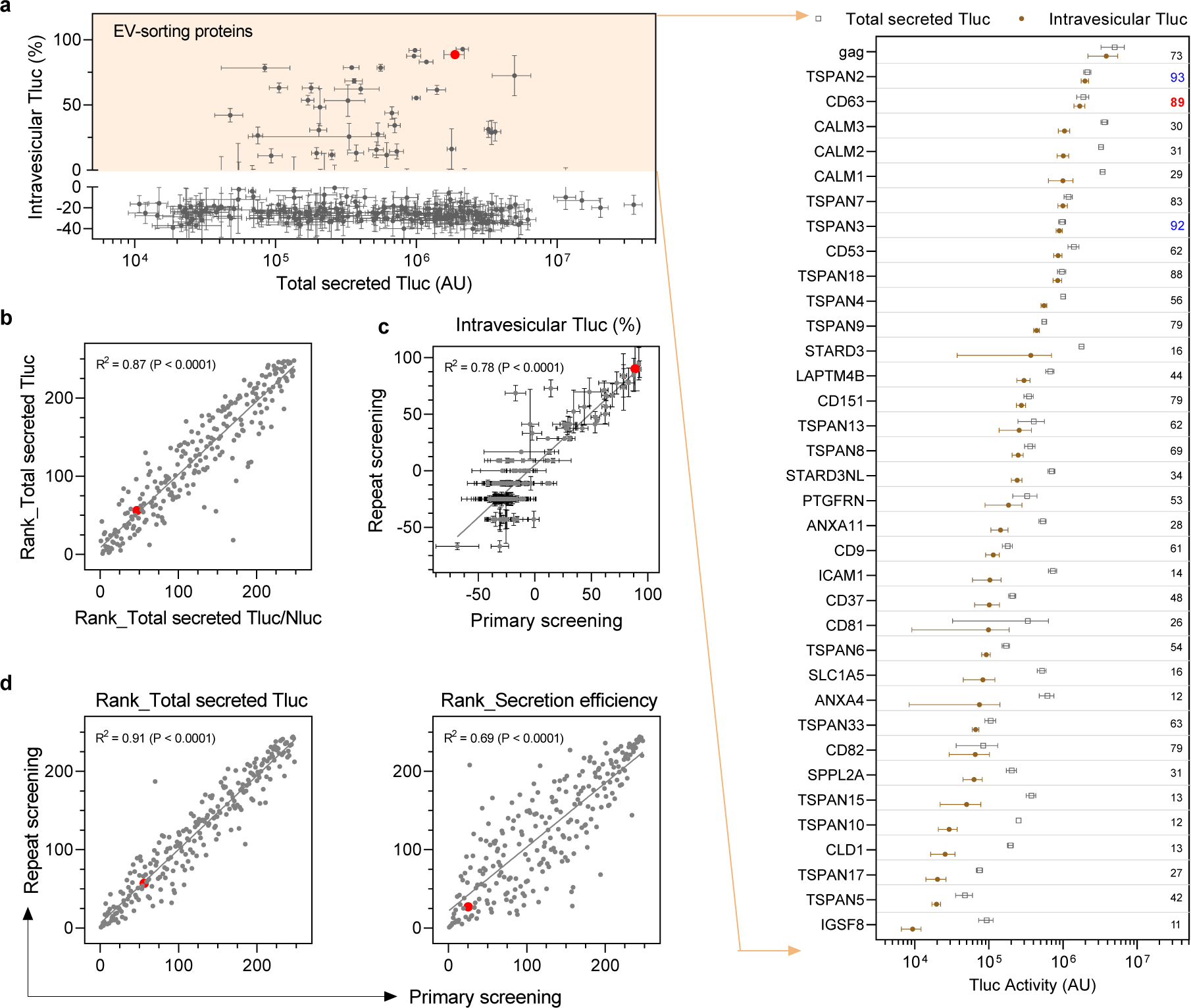
EV-sorting ability of the candidate proteins. (a) Overview of all 244 candidates plotting total secreted Tluc against percentage of intravesicular Tluc. EV-sorting proteins were defined to have a percentage of intravesicular Tluc above zero (yellow area), which are all shown in the grouped dot plot. The value refers to the percentage of intravesicular Tluc to total secreted Tluc. Proteins are marked with gene names. (b) Correlation between the rank of total secreted Tluc and the rank of total secreted Tluc/Nluc ratio. (c) Correlation of the percentage of intravesicular Tluc obtained from the primary and repeat screening. (d) Correlation of the rank of secreted Tluc or secretion efficiency between the primary and repeat screening. Results in (a-b) were from the primary screening and are shown as mean ± standard deviation of five biological replicates. Data from the repeat screening are shown as mean ± standard deviation of three biological replicates. In the scatter plots, each dot refers to one candidate and the red dot indicates the benchmark CD63. The degree of correlation was analyzed with linear regression and is shown as goodness-of-fit (R^2^) and significance of non-zero slope (P).

Out of the four known non-viral EV-sorting proteins, CD63 showed the highest percentage of intravesicular Tluc (89%). with 89% of total secreted Tluc localized inside EVs. Strikingly, TSPAN2 outperformed CD63 not only in terms of relative intravesicular Tluc (93%) but also absolute amount (by 18%; Table S2). Apart from that, three calmodulin proteins (CALM1, CALM2, CALM3) sorted considerable amounts of absolute Tluc into EVs but with moderate percentages of intravesicular Tluc (29-31%).

To rule out that any unwanted factors interfered with Tluc secretion, suitable quality controls were put in place. First, all candidates were ranked according to total secreted Tluc and the ratio of total secreted Tluc to Nluc. Normalization against Nluc signal did not affect Tluc secretion (Figure 2b), thereby dismissing a confounding role of the transfection procedure. Secondly, a repeat of the screening revealed consistent results for the percentage of intravesicular Tluc (Figure 2c) as well as the ranks of total secreted Tluc and secretion efficiency (Figure 2d).

Finally, the ranks of total secreted Tluc and secretion efficiency showed a high degree of linear correlation between two different plasmid doses (0.75 µg/mL vs 1 µg/mL; Figure S1c). Taken together, these results substantiate the reliability of the findings obtained with our screening method.

### EV-sorting proteins are largely conserved across different cell types

Besides HEK-293T, other cell types are regularly used as EV sources prompting us to screen the EV-sorting ability of our candidates in (1) suspension-adapted HEK cells (Freestyle 293-F), (2) human cord blood-derived mesenchymal stem cells (MSCs), (3) human hepatocyte-derived carcinoma cells (Huh-7), and (4) mouse kidney epithelial cells (TCMK-1). For this purpose, 95 candidates that had shown promise (top 40) in the initial screening in either of these categories: percentage of intravesicular Tluc, total secreted Tluc, and secretion efficiency (Table S2), were screened as above.

In Freestyle 293-F the protein with the highest EV-sorting ability in terms of percentage of intravesicular Tluc was TSPAN3. In MSCs and Huh-7 CALM2 occupied the highest rank, and in TCMK-1 CD63 (Figure 3a). While the transfection procedure was not found to substantially affect Tluc secretion in all the adherent cells tested, Freestyle 293-F seemed more prone to variation (Figure S2). Between 30 and 37 proteins with EV-sorting ability (percentage of intravesicular Tluc above zero) were identified for each cell type (Table S3). Out of these, 24 proteins were conserved across all five cell types (Figure 3b) indicating their robust sorting ability in different cellular contexts. The proteins in the conserved subset were ranked according to their absolute intravesicular Tluc activity (Figure 3c). On average, TSPAN2, CD63 and TSPAN3 demonstrated the best sorting ability across different cell types.

**Figure 3.**
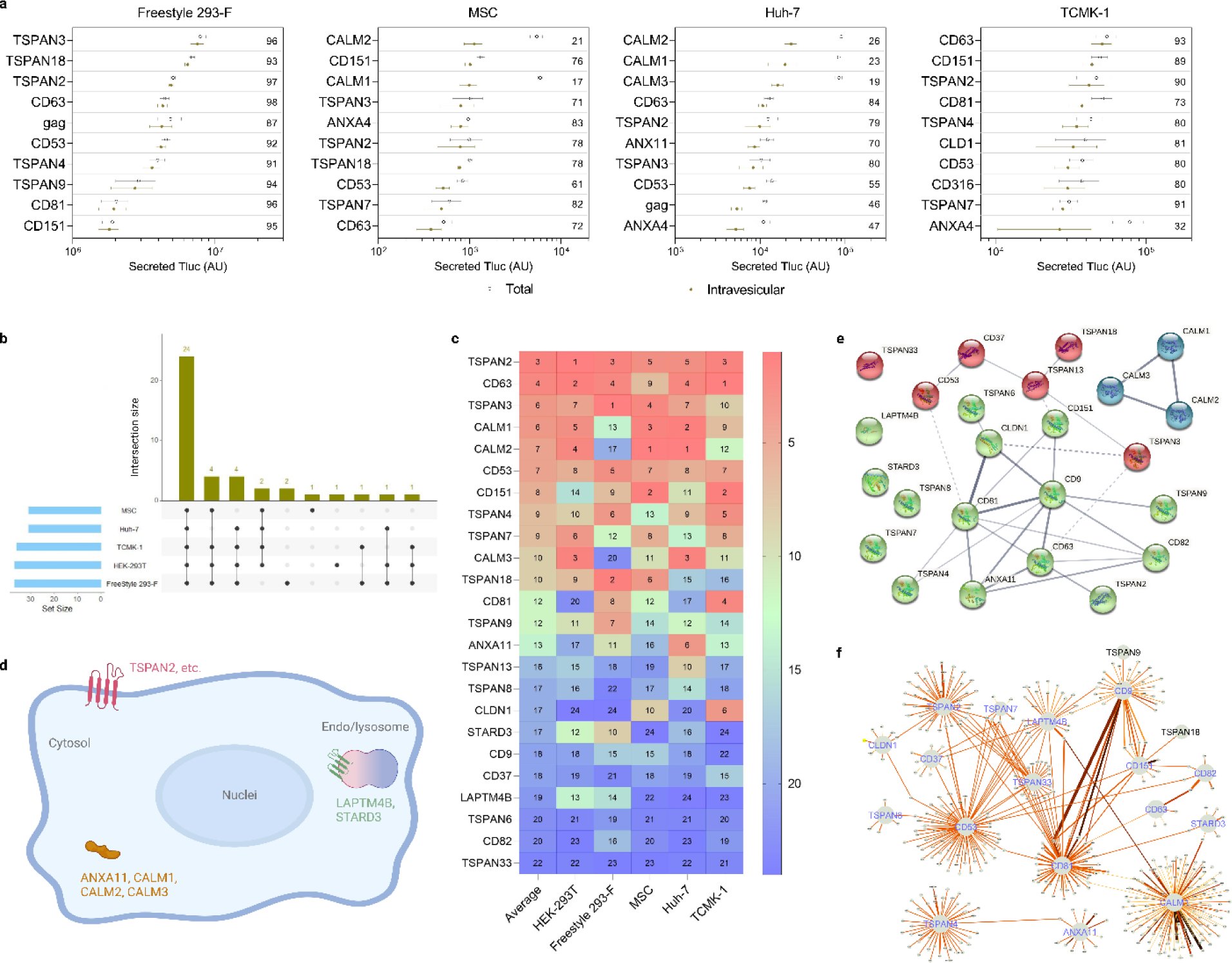
EV-sorting ability in various cell sources. (a) The top ten scaffold proteins regarding intravesicular Tluc in different producer cell types. The value inside the plot refers to the percentage of intravesicular Tluc. Results are shown as mean ± standard deviation of three biological replicates. (b) Number of EV-sorting proteins identified for each producer cell type and overlap between cell types. (c) Ranking the 24 conserved EV-sorting proteins according to their absolute intravesicular Tluc activity in each cell type. The number indicates the rank in each cell type as well as the average thereof. (d) Topology and subcellular location of the 24 conserved EV-sorting proteins. (e-f) Interaction network of the 24 conserved EV-sorting proteins retrieved from STRING (e) and IntAct (f) databases. Line thickness in panel (e-f) indicates the strength of data support. Proteins are marked with gene names.

To obtain a more in-depth understanding of potential mechanisms that govern the EV-sorting ability of the conserved subset, bioinformatic studies were conducted. According to the annotation available on UniProtKB, only the three calmodulin proteins and ANXA11 are cytosolic, while the remaining proteins are all members of the tetraspanin superfamily and located either on the plasma membrane or that of endosomes/lysosomes (Figure 3d). Next, we evaluated possible interactions between the 24 EV-sorting proteins. The experimental and predicted interactome available from the STRING database showed weak evidence for the interaction of TSPAN2/TSPAN3 with CD63, and calmodulin proteins seem to operate irrespective of the rest (Figure 3e). Similarly, the results from the IntAct database, which includes both direct and indirect interactions, suggested that the interactome of TSPAN2 overlaps poorly with that of the three well-characterized tetraspanins CD9/CD63/CD81 (Figure 3f). These predictions indicate that TSPAN2 and TSPAN3 operate largely independent of each other and other tetraspanins.

### EV-sorting candidates prove robust amid standardized EV production

In the screening, HEK-293T cells were grown and transfected in 96-well microplates for the purpose of higher throughput. Such a scale, however, is not practicable for future applications seeking to produce larger quantities of engineered EVs. Additionally, the conditioned media in the screening were analyzed directly after centrifugation without any defined separation techniques. Here, we were particularly interested in small EVs (sEVs, ≤ 200 nm) because of their therapeutic potential in many diseases^36, 37^. With that in mind, we produced EVs according to a standardized protocol recently established by our group^38^. The main differences to the initial screening protocol were (1) a higher dose of plasmids and shorter transfection duration, (2) subsequent maintenance in Opti-MEM, and (3) filtration of conditioned media through a 200-nm membrane followed by a concentration step (Figure 4a). Additionally, sEVs were separated from soluble proteins by SEC before measuring Tluc activity and vesicle counts (Figure 4b). Noteworthily, Tluc in the eluate was completely deactivated by Proteinase K indicating that resistant protein aggregates were not a matter of concern (Figure 4b).

**Figure 4.**
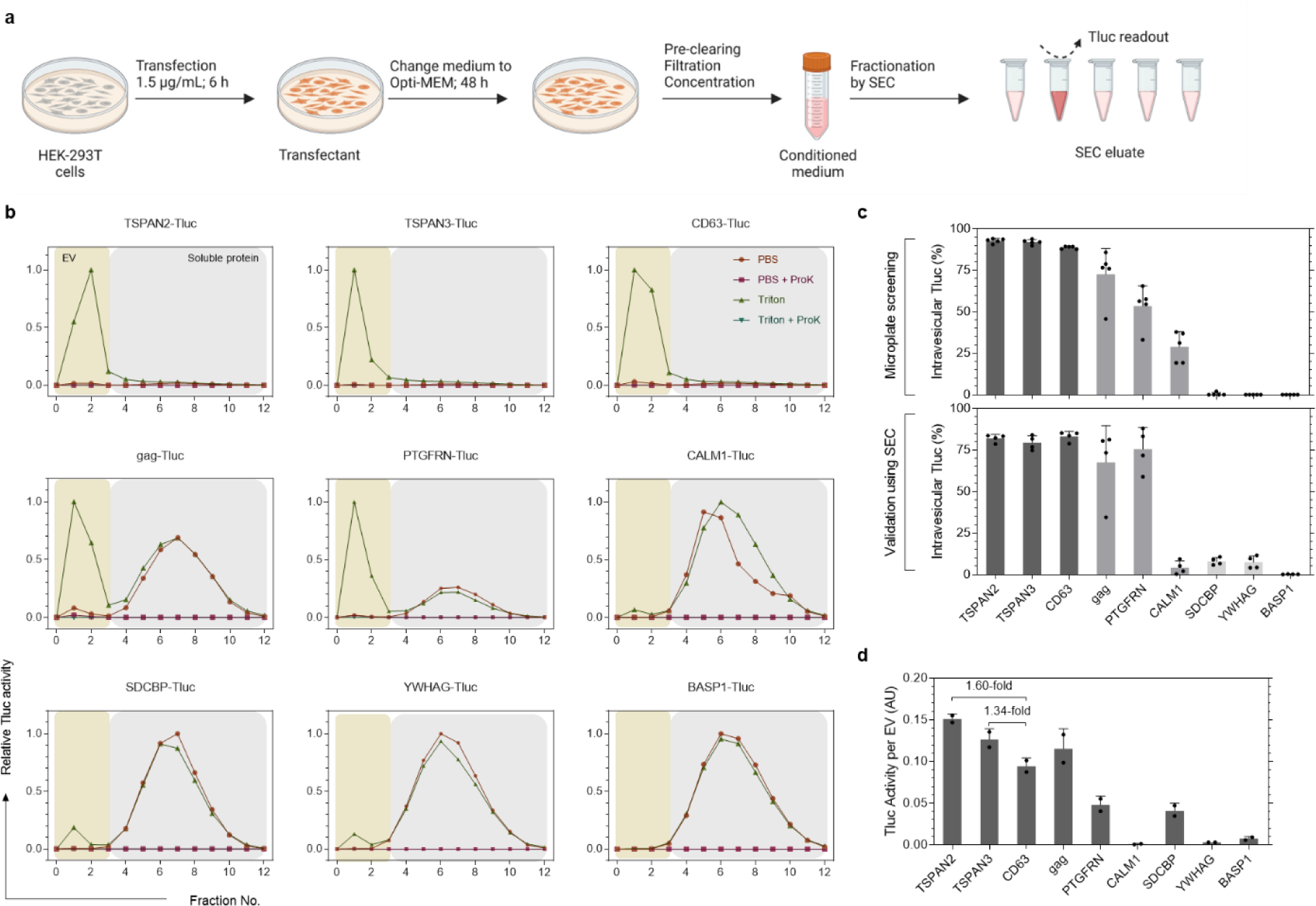
EV-sorting ability of candidate proteins in standardized production conditions. (a) Workflow of the EV production protocol and SEC fractionation. (b) Tluc-based SEC elution profiles of conditioned media from transfected HEK-293T cells. EVs and soluble proteins were recovered in fractions 0-3 and 4-12, respectively. Tluc activity in each fraction was measured with and without Triton and ProK. Data was normalized to the fraction with the highest signal. (c) Percentage of intravesicular Tluc from the screening (upper panel; five biological replicates) and standardized (lower panel; four biological replicates) protocols. (d) Calculated Tluc activity per vesicle for purified EV preparations. Results are shown as mean ± standard deviation of two biological replicates. Proteins are marked with gene names.

To gain insight into sEV-sorting ability, nine representative candidates were selected based on their performance in the screening (Figure 4c). The percentage of intravesicular Tluc generally coincided with the screening, showing high (> 80%; TSPAN2, TSPAN3, and CD63) and low (< 15%; SDCBP, YWHAG, BASP1) sEV-sorting ability (Figure 4c). Interestingly, CALM1 sorted only 3.7% Tluc into sEVs as compared to 28.9% in the screening (P < 0.01, student T test).

Taking vesicle numbers into account, we observed that TSPAN2 and TSPAN3 outperformed CD63 in terms of Tluc activity per EV by 60% and 34%, respectively (Figure 4d). Overall, these results demonstrate that the sorting ability of these nine candidates remained largely unchanged when following a standardized sEV production protocol.

### EV-sorting candidates prove versatile for different cargos

Luciferase is a facile reporter for quantifying engineered EVs in bulk. However, to obtain more information on the number of engineered EVs and the abundance of cargo proteins per EV, single-vesicle imaging flow cytometry is the method of choice^39, 40^. Therefore, for the nine candidate proteins explored above, Tluc was replaced with a hybrid reporter consisting of the fluorescent protein mNeonGreen (mNG; 26.6 kDa) fused to HiBiT^41^, an 11-mer peptide from split Nluc luciferase (Figure 5a).

**Figure 5.**
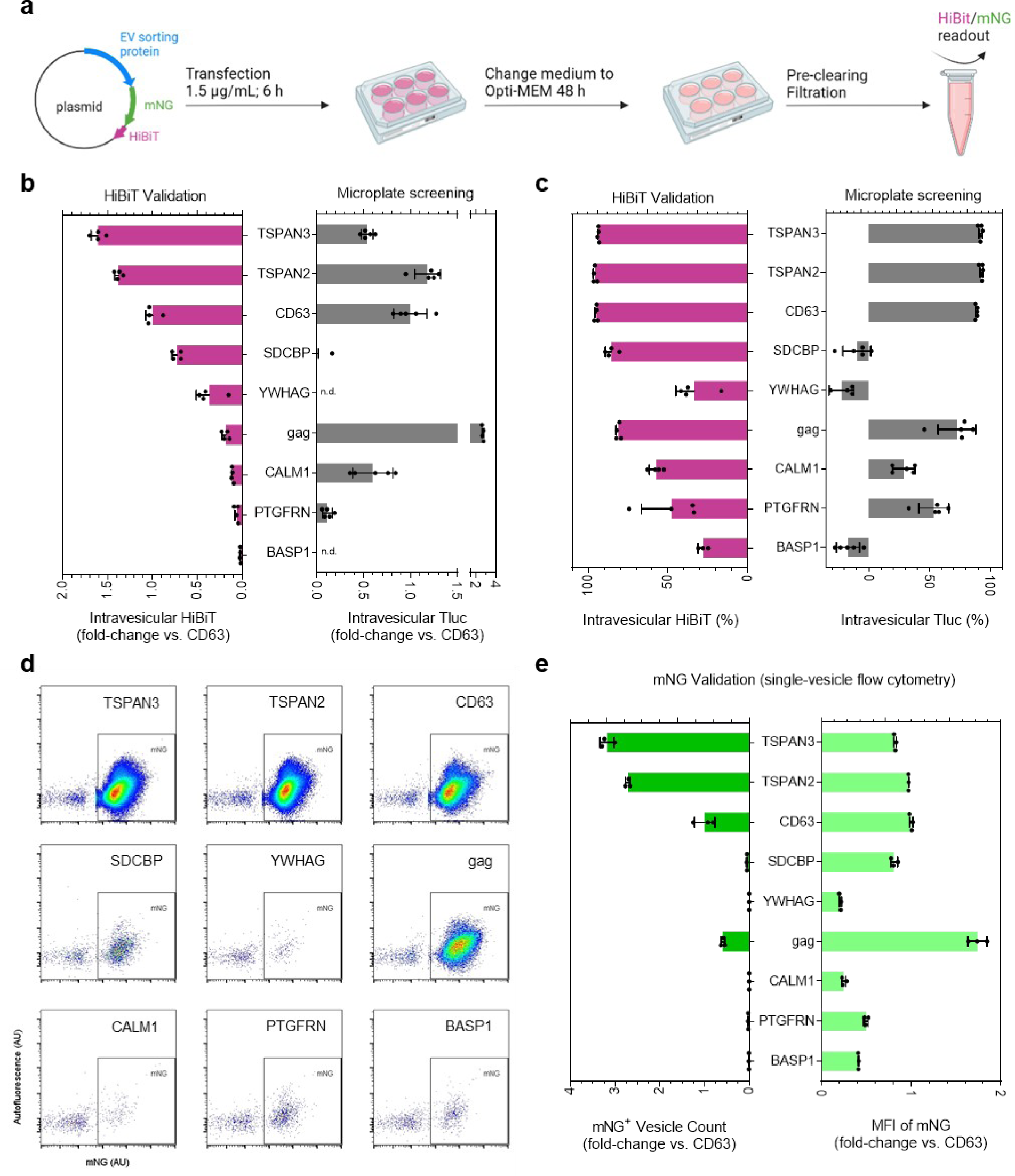
EV-sorting ability of candidate proteins fused to a hybrid bioluminescent and fluorescent reporter. (a) EV production and analysis workflow. (b) Intravesicular HiBiT relative to the benchmark CD63. n.d.: not detected. (c) Percentage of intravesicular HiBiT. In (b-c), results are shown as mean ± standard deviation of four biological replicates. Screening results from HEK-293T cells were graphed for reference. (d) Single-vesicle flow cytometry dot plots of mNG-HiBiT-labeled EVs. (e) Concentration and mean fluorescence intensity (MFI) of mNG-positive EVs. Results are shown as mean ± standard deviation of three biological replicates. Proteins are marked with gene names.

Comparison of intravesicular luciferase activities of HiBiT and Tluc revealed comparable engineering efficiencies for TSPAN3, TSPAN2 and CD63 in terms of amount (Figure 5b) and percentage (Figure 5c). Using mNG to look at the single-vesicle level, TSPAN3 and TSPAN2 produced the highest number of engineered EVs, outperforming CD63 by around 3-fold while reaching similar levels of mNG per engineered EV (Figure 5d-e). Additionally, we showed that mNG co-localized with respective sorting proteins on the EVs after antibody staining, which is indicative of intact fusion proteins (Figure S3). Surprisingly, HiBiT- and Tluc-based measurements differed greatly for gag (Figure 5b-c); however, its mNG levels did not show such a discrepancy (Figure 5d-e). This led us to postulate that the steric configuration of gag-mNG-HiBiT prohibits HiBiT from complexing with its partner subunit to form functional luciferase. Additionally, CALM1 showed low levels of vesicular HiBiT and mNG, in line with the trend observed for CALM1-Tluc in SEC validation experiments (Figure 4c). These results suggest that CALM1 preferentially sorts into larger vesicles (>200 nm) that are removed during the filtration step (Figure S4).

We additionally tested the performance of these selected proteins in Freestyle 293-F cells, which are a prominent source for EV production due to a less tedious propagation procedure. Again, TSPAN3 and TSPAN2 outperformed CD63 by 82% and 50%, respectively, in terms of intravesicular HiBiT (Figure S5a). On a single-vesicle level, the highest concentration of mNG-positive vesicles was produced by TSPAN3 and TSPAN2 engineered Freestyle 293-F cells (Figure S5b). Collectively, the reconciling results based on mNG-HiBiT and Tluc reporters reinforce the reliability of the screening protocol and highlight the robust EV-sorting ability of the candidate proteins for different cargos.

### Distinct molecular signatures among tetraspanin-engineered EVs

Throughout all experiments, TSPAN2 and TSPAN3 appeared among the best EV-sorting proteins, seemingly performing better than the well-characterized tetraspanin CD63. Notably, different splice isoforms of TSPAN2 and TSPAN3 failed to retain EV-sorting ability in HEK293-T cells (Figure S6). Since both proteins are new to the EV field, we aimed to characterize the physiochemical features of TSPAN2- and TSPAN3-engineered EVs in relation to CD63-engineered EVs in greater detail.

Nanoparticle tracking analysis of EV preparations from transfected HEK-293T cells revealed a narrow size distribution with median hydrodynamic diameters of approximately 120 nm (Figure 6a). Also, their morphological appearances were typical of EVs as exemplified by membrane structure and size (Figure 6b). Moreover, common EV markers such as CD81, syntenin-1 and TSG101, but not the negative marker Calnexin, were detected in the EVs (Figure 6c). Apart from that, we examined the location of the three tetraspanin proteins in transfected cells to gain insight into EV biogenesis. While TSPAN2 was localized in both the plasma membrane and cytosol of producer cells, TSPAN3 and CD63 were primarily detected as punctate signals inside cells (Figure 6d). Furthermore, the production of all types of engineered EVs was resistant to inhibition of ceramide (Figure S7a), which is a driver of a recognized exosome production pathway^1, 42^.

**Figure 6.**
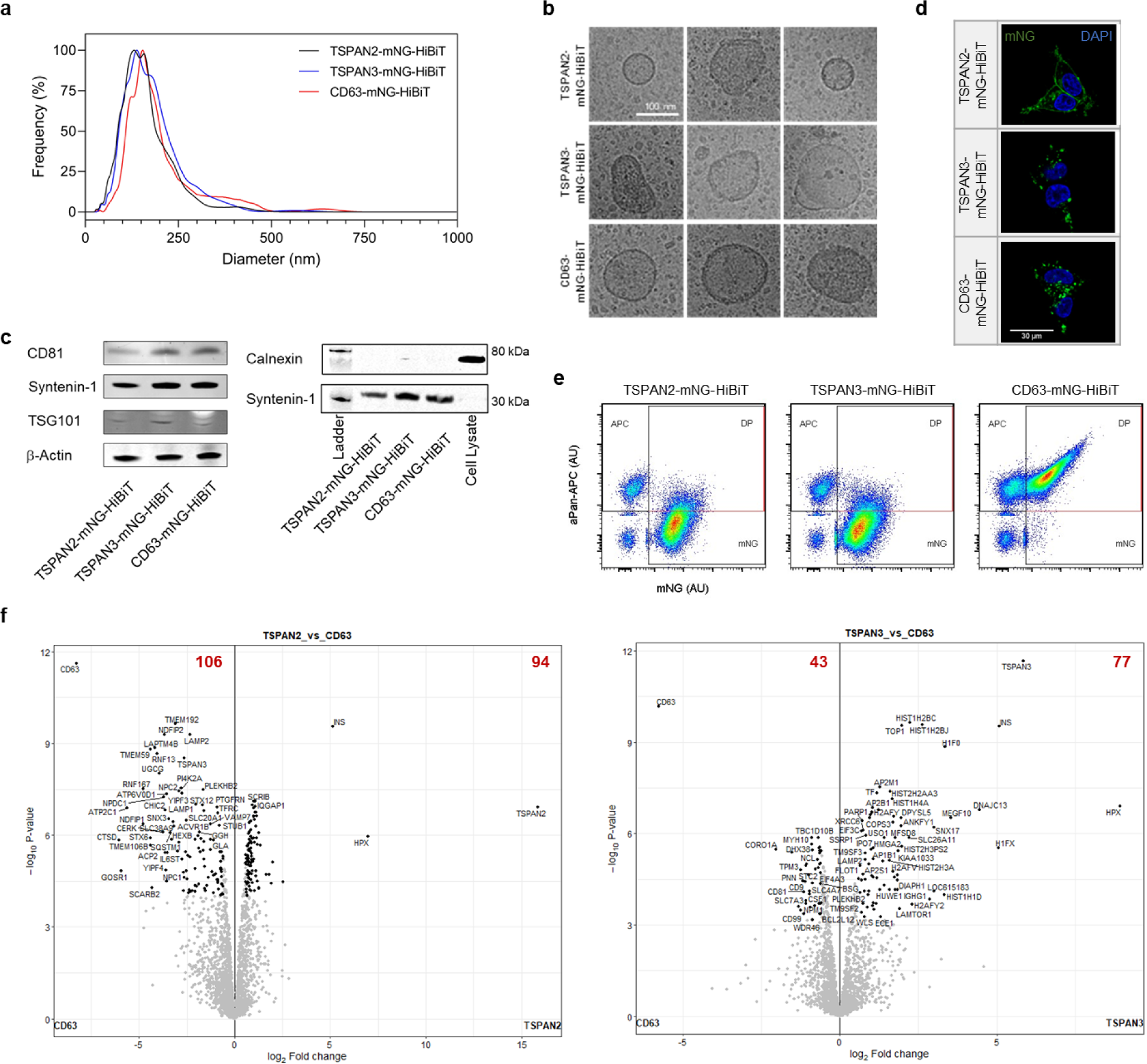
Physiochemical features of TSPAN2-, TSPAN3- and CD63-enriched EVs. (a) Size distribution of EVs from transfected HEK-293T cells. (b) Representative cryo-electron microscopy images of EVs. (c) Western blots of common markers and exclusion markers of EVs. (d) Cellular location of tetraspanins in transfected HEK-293T cells. (e) Single-vesicle flow cytometry dot plots of EVs after staining with APC-conjugated tetraspanin antibodies. Pan refers to CD63/CD81/CD9. (f) Volcano plots showing differentially regulated proteins in EVs. Results were from three biological replicates.

Furthermore, to get an understanding of their protein signatures, EVs were stained with the classic pan-surface markers CD9/CD63/CD81 and analyzed on a single-vesicle level. Strikingly, only a small fraction of TSPAN2- and TSPAN3-engineered EVs displayed the three classic EV markers on their surface (Figure 6e). In addition, EV surface expression analysis of 39 proteins by multiplex bead-based flow cytometry^43^ revealed that the surface epitope composition of TSPAN2-positive EVs differed from that of CD9/CD63/CD81-positive EVs (Figure S7b).

Besides surface proteins, in-depth proteomic analyses of EVs showed a plethora of differentially expressed proteins for TSPAN2 (106 downregulated and 94 upregulated) and TSPAN3 (43 downregulated and 77 upregulated) compared to CD63-engineered EVs. CD9/CD63/CD81 were among the proteins that were significantly downregulated in TSPAN2/TSPAN3-engineered EVs, which is in line with the results obtained from single-vesicle and bead-based flow cytometry.

Based on Gene Ontology clustering, in comparison with wildtype EVs from HEK293T cells, all three types of engineered EVs (CD63, TSPAN2 and TSPAN3) were enriched (> 10%) with metabolite interconversion enzymes, protein modifying enzymes, and RNA metabolism proteins, but depleted (> 10%) of extracellular matrix proteins (Figure S8). The differences were also apparent in overall protein composition, where engineered EVs clustered away from wildtype EVs and to a lesser extent also from each other (Figure S7c). Interestingly, we observed that TSPAN2 and TSPAN3 are only present in EVs upon over-expression (Figure S7d). Moreover, in relative to WT EVs, overexpression of TSPAN2 negatively impacted CD63 and TSPAN3 levels, which could indicate a competitive relationship. TSPAN3 overexpression, on the other hand, led to a slight increase in CD63 levels suggesting a positive regulation (Figure S7d). Overall, these findings illustrate that TSPAN2/TSPAN3-based engineering gives rise to EV subpopulations distinct from CD63.

### TSPAN2- and TSPAN3-engineered EVs as delivery modalities

To explore whether TSPAN2/TSPAN3-engineered EVs are suitable for cellular delivery, we investigated their delivery potential *in vitro* and *in vivo*. First, Huh-7 cells were treated with mNG-labeled EVs to examine their subcellular location in recipient cells. The strong punctate yellow signal clearly indicated efficient internalization and trafficking to lysosomes (Figure 7a). Quantification of cellular MFI using flow cytometry revealed slightly better uptake efficiencies for TSPAN2/TSPAN3-engineered EVs compared to CD63-engineered EVs (Figure 7b). For EV distribution studies in mice, equal amounts of engineered EVs (based on Tluc activity) were administered intravenously and tracked in real time with in vivo imaging system (Figure S9a, Figure 7c). For all three types of engineered EVs, we observed rapid distribution to liver and spleen within five minutes (Figure 7c) and a notable decline in whole-body activity over 30 min (Figure S9c, P = 0.0006, Kruskal-Wallis test). TSPAN2- and TSPAN3-engineered EVs seemed to confer slightly higher whole-body retention than CD63-engineered EVs (Figure S9b). Results from subsequent *ex vivo* measurements supported the dominant hepatic and splenic accumulation of engineered EVs (Figure 7d). Taken together, similar to CD63-engineered EVs, TSPAN2- and TSPAN3-engineered EVs are efficiently taken up by cells *in vitro* and *in vivo*.

**Figure 7.**
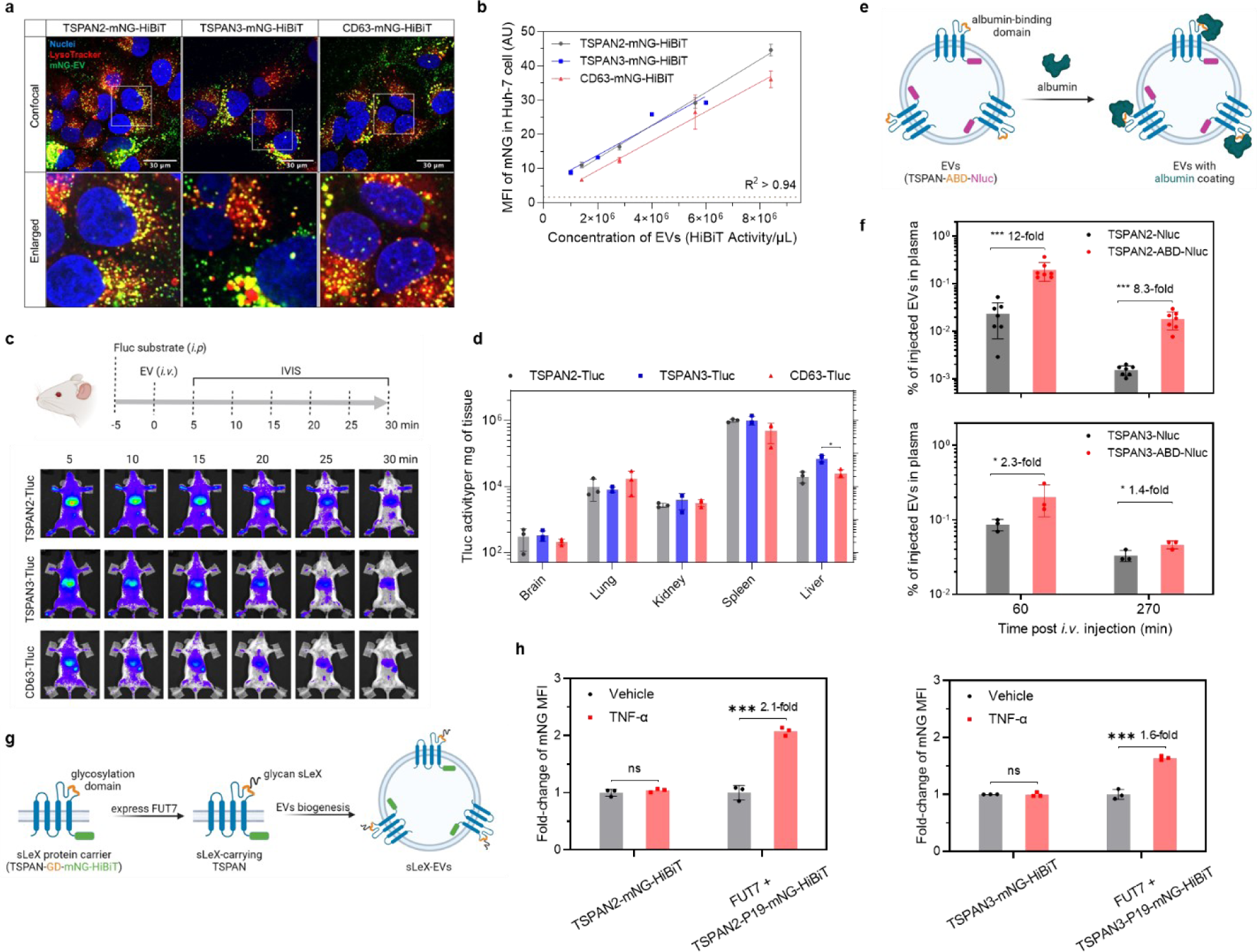
Biological activity of TSPAN2- and TSPAN3-engineered EVs. (a) Huh-7 cells were treated with mNG-HiBiT-labeled EVs for 4 h and stained with LysoTracker to visualize lysosomes. Representative confocal microscopy images are shown. (b) Huh-7 cells were treated with mNG-HiBiT-labeled EVs for 8 h. Cellular mNG MFI was quantified using flow cytometry. Data are shown as mean ± standard deviation of three biological replicates. The degree of correlation was analyzed with linear regression and is shown as goodness-of-fit (R^2^). (c) Biodistribution of Tluc-labeled EVs in mice. NMRI mice were intraperitoneally (*i.p.*) injected with D-luciferin substrate. Five minutes later, mice were intravenously (*i.v.*) injected with the same amount of engineered EVs (based on Tluc activity) and imaged with IVIS. Subsequently, major organs were collected for *ex vivo* bioluminescence measurements. Representative IVIS images are shown. (d) Quantification of Tluc activity in organs *ex vivo* after IVIS. Results are shown as mean ± standard deviation of three mice. (e) Scheme of tetraspanin engineering for generating albumin-binding EVs. EVs were collected from HEK-293T cells stably expressing the fusion proteins. (f) Albumin-binding EVs were injected intravenously and their concentration in plasma was determined. Data are shown as mean ± standard deviation of 3-7 mice. (g) Scheme of tetraspanin engineering for sLeX display on EVs. EVs were collected from HEK-293T cells stably expressing the components. (h) HUVEC cells were activated by TNF-α for 2 h and treated with EVs for 6 h. Cellular mNG MFI was quantified using flow cytometry and is shown as fold-change over un-activated cells. Data are shown as mean ± standard deviation of three biological replicates. Two-tailed Student T test. ns: not significant; *: P ≤ 0.05; ***: P ≤ 0.001.

Having extensively showcased the loading and delivery capacity of TSPAN2 and TSPAN3 for luminal cargo, we next investigated their potential for EV surface display applications. The large extracellular loops (LELs) of some tetraspanins have already been exploited for such applications and given the topological similarities of tetraspanin proteins, we sought to engineer the LELs of TSPAN2 and TSPAN3. Insertion of an albumin-binding domain (ABD) into the LEL of CD63, CD9, and CD81 has been shown to extend the plasma circulation time of EVs drastically^44^. Using the same strategy, an ABD was cloned into the LEL of TSPAN2 and TSPAN3 with Nluc at the C-terminus for quantification purposes (Figure 7e). EVs were collected from HEK-293T cells stably expressing TSPAN-ABD-Nluc fusion proteins (Figure S9c) and assessed for their albumin-binding ability. As expected, only ABD-displaying EVs bound to albumin (Figure S9d). Next, these EVs were intravenously injected into mice and EV concentrations in plasma were measured on the basis of Nluc at different timepoints. In comparison to wildtype tetraspanin-engineered EVs, ABD-displaying EVs had significantly higher concentration in plasma, particularly when using TSPAN2 as the scaffold protein (Figure 7f).

In another example, we focused on cell-specific targeting with glycan ligand sialyl Lewis X (sLeX) for specificity towards activated endothelial cells^45^. Here, a 19-mer sLeX peptide carrier (P19) was inserted into the LEL of each tetraspanin protein with mNG-HiBiT at the C-terminus. In the presence of fucosyltransferase VII (FUT7), P19 is glycosylated to display sLeX (Figure 7g). Based on this rationale, sLeX-EVs were produced from HEK-293T cells stably expressing FUT7 and TSPAN-P19-mNG-HiBiT (Figure S9c). Their uptake was evaluated in TNF-a-activated endothelial cells, which express E-selectin, the main receptor for sLeX. Wildtype tetraspanin-engineered EVs were taken up similarly in un-activated and activated endothelial cells while sLeX-EVs, using either TSPAN2 or TSPAN3 as the scaffold, demonstrated preferable uptake by activated endothelial cells (Figure 7h). Overall, this demonstrates the feasibility of simultaneous engineering of the LEL and C-terminus of TSPAN2 and TSPAN3 for surface display and luminal cargo loading, thus highlighting their potential for therapeutic applications.

## DISCUSSION

In this study, we successfully established a simple and robust assay to quantify the EV-sorting ability of proteins, which is instrumental for endogenous cargo loading into the lumen of EVs. In doing so, we were the first to identify and qualify TSPAN2 and TSPAN3 as highly efficient EV-sorting candidates. Both TSPAN2 and TSPAN3 consistently appeared among the top-performing candidates across different cell lines and for different cargo proteins (sizes ranging from 26.6 kDa to 60.5 kDa). Furthermore, the cellular uptake of TSPAN2- and TSPAN3-engineered EVs *in vitro* and *in vivo* demonstrates their great potential as improved delivery modalities. The fact that neither TSPAN2 nor TSPAN3 are highly expressed in EVs from wildtype HEK-293T cells (Figure S7d) might explain why they had been neglected in previous EV engineering studies.

Consequently, little is known about their functions especially in relation to EVs. TSPAN2 is hypothesized to be involved in oligodendrocyte differentiation and cancer metastasis^46^. Less is known about the functions of TSPAN3 besides a link to the progression of acute myeloid leukemia^47^, however, its presence in urinary exosomes has been shown^48^. Nevertheless, our functional characterization of TSPAN2- and TSPAN3-engineered EVs suggests similar internalization dynamics and fate as CD63 *in vitro* and *in vivo*.

The majority of candidates (21 out of 24) with conserved EV-sorting ability across different cell types was found to belong to the tetraspanin superfamily, demonstrating the importance of this topological feature for EV sorting. While tetraspanins are implicated in a variety of cellular processes, some of them have been found to aid membrane curvature therefore implying a role in EV biogenesis^49, 50^. By inhibiting ceramide-dependent EV biogenesis, we showed that neither TSPAN2-, TSPAN3-nor CD63-engineered EVs are produced via this route. Other studies suggest an ESCRT-independent pathway for CD63^51, 52^, which altogether supports the concept of a tetraspanin-dependent route of EV biogenesis^53^. Consequently, overexpression of tetraspanins could boost the production of (engineered) EVs. However, some family members might be more useful to that end than others, which is reflected in their heterogenous EV-engineering ability observed here. This is also connected to our and others’ observations that distinct tetraspanin subpopulations of EVs exist^27, 54^. Hence, we hypothesize that overexpression of tetraspanins results in the production of certain EV subpopulations, thereby dictating its EV-engineering potential. However, unraveling the individual implications of tetraspanins in EV biogenesis is a matter of concern for future studies.

Interestingly, we found that calmodulins preferentially sort cargo into larger EVs (>200 nm). Given that larger EVs are commonly generated by outward budding of the plasma membrane, we believe that calmodulins are predominantly present in microvesicles (100-1000 nm). In support of that, CALM1 showed weak putative interactions with typical markers of small EVs produced in the endosomal system, i.e., exosomes. This finding can prove particularly useful to applications involving large EVs for drug or gene delivery.

Although we aimed to address various aspects of endogenous EV engineering, future studies will provide further insight into the EV-sorting ability of our candidates in other settings. Cargo proteins here were fused to the C termini of all 244 proteins, leaving no interpretation for other fusion sites. Despite that our screening was designed to compare luminal cargo proteins of different EV-sorting candidates, we showed that the lead proteins TSPAN2 and TSPAN3 were tolerable to simultaneous surface engineering and luminal cargo loading. Additionally, it remains to be determined whether the EV-sorting ability observed here upholds for alternative cargo molecules. These or other reasons could potentially explain why we were unable to observe superior EV-sorting ability for BASP1 and PTGFRN as seen in a previous study^32^. Apart from that, it is not surprising to observe producer cell-dependent EV sorting ability. HIV-derived gag, for example, displayed impressive sorting ability in HEK-293T cells but not in MSCs (Table S3). Other candidates were found to be exclusive to certain cell types likely influenced by the cell’s transcriptome and proteome, therefore demanding individual screenings. On the upside, a conserved subset of proteins was identified that we hypothesize to be involved in core physiological processes of EV production. These candidates lend themselves as reliable EV-engineering candidates irrespective of the producing cell.

In conclusion, this study is by far the most comprehensive of its kind that examined the EV-sorting potential of overexpressed proteins. Hence, it will provide a valuable reference point for researchers aiming to sort cargos into EVs by endogenous means. Additionally, TSPAN2 and TSPAN3 are first to be identified here as reliable and efficient EV-sorting proteins, which poses a new steppingstone for endogenous engineering strategies and might in foresight broaden the applications of EVs as delivery modalities.

## METHODS

### Cloning

Codon-optimized DNA sequences coding for the scaffold protein and luciferase reporter were cloned downstream of the CAG promoter into the pLEX vector (Twist Bioscience, US). ABD and P19 peptides were inserted between amino acid (aa) 154-155 for TSPAN2 and aa 150-151 for TSPAN3, respectively. To generate different constructs expressing mNG-HiBiT or Nluc, protein-coding sequences for Tluc were replaced with corresponding fragments through In-Fusion cloning (Takara; 638948) or restriction cloning strategies. All expression cassettes were confirmed by sequencing. Scaffold protein identifiers are listed in Table S1. Plasmids are available from the corresponding author upon request.

### Cell culture

HEK-293T, Huh-7 and TCMK-1 cells were maintained in high glucose DMEM media (Gibco, 41966-029) supplemented with 10% fetal bovine serum (FBS; Gibco, 10270-106) and 1% anti-anti (Gibco, 15240). HUVEC cells were cultured in Endothelial Cell Growth Medium MV 2 (PromoCell, C-22022) supplemented with 1% anti-anti. Human cord blood-derived MSCs were cultured in MEM media (Gibco, 22561-021) supplemented with 10% FBS and 1% anti-anti. Freestyle 293-F cell were kept in FreeStyle 293 Expression Media (12338-018) under continuous shaking at 175 rpm. All cells were cultured in humidified incubators with 37°C and 5% CO_2_.

### Transfection

In the screening protocol, producer cells (HEK-293T, Huh-7 and TCMK-1) were seeded in 96-well plates (100 µL media per well) and transfected when approximately 30% confluent. Freestyle 293-F cells were seeded at 7.5×10^4^ cells per well (90 µL media per well) and directly transfected. For transfection, 10 µL of plasmid-lipofectamine mixture containing 75 ng plasmid and 165 ng Lipofectamine 2000 (Invitrogen, 11668-019) in Opti-MEM (Gibco, 31987-047) were added to each well. The cells were cultured for an additional 48 hours before the conditioned media were collected.

In 6-well plates, HEK-293T cells were seeded in 2 mL media per well and transfected when approximately 60% confluent. 3 µg of plasmid were complexed with 6.6 µg of Lipofectamine 2000 in 200 µL Opti-MEM and added to each well. The cells were cultured for an additional 48 hours before the conditioned media were collected.

In 15-cm petri dishes, HEK-293T were seeded in 20 mL media per petri dish and transfected when approximately 60% confluent. 30 µg of plasmid were complexed with 45 µg of polyethyleneimine (Polysciences; 24765-1) in 4 mL Opti-MEM and added to each dish. Transfection was discontinued 6 h later by changing the media to Opti-MEM and the cell culture was maintained for an additional 48 h before harvesting conditioned media.

### Establishing stable cell lines

Lentivirus encoding transgenes of interest were produced in HEK-293T cells according to our previous reports.^55, 56^ To generate stable cell lines, HEK-293T cells were cultured in 6-well plates until approximately 60% confluent and then transduced with lentiviral particles overnight. Transduced cells were expanded and selected using 4 µg/mL of puromycin (Sigma-Aldrich, P8833).

### Isolation of extracellular vesicles

In the screening protocol, conditioned media from transfected cells was pre-cleared by two rounds of centrifugation (700×g for 5 min and then 2000×g for 10 min) to pellet cells and cell debris. If not specified, the supernatant was filtered through 200 nm membrane to remove large particles. To obtain enough EVs for size exclusion chromatography (SEC), 20 mL of the processed media was concentrated to approximately 1 mL using Amicon Ultra-2 spin-filter with 10 kDa molecular wight cut-off (Millipore, UFC201024).

To produce EVs in larger scale, we followed a protocol previously reported by our group^38^. Briefly, after pre-clearing and filtration (s.a.), large volumes of conditioned media were diafiltrated and concentrated to roughly 50 mL using the KrosFlo KR2i TFF System (Repligen, US) with 300 kDa cut-off hollow fiber filters (Spetrum Labs, D06-E300-05-N) at a flow rate of 100 mL/min (transmembrane pressure at 3.0 psi and shear rate at 3700 sec^−1^)^57^. EVs were further concentrated until approximately 500 μL using Amicon Ultra-15 spin-filter with 100 kDa molecular weight cut-off (Millipore, UFC910024) and stored at −80 °C in PBS-HAT buffer^58^ before downstream analysis.

### Size exclusion chromatography

500 µL of concentrated conditioned media was loaded onto SEC columns (Izon, SP1). After discarding the void fraction (first 2 mL), a total of 13 fractions of the eluate (1 mL per fraction) was collected sequentially and numbered as 0-12. According to the manufacturer’s instruction, fraction 1-3 were enriched with EVs and pooled as the EV fraction for downstream analysis. For better separation, the eluate was collected into 48 fractions (0.3 mL per fraction). To check albumin-binding ability *in vitro*, EVs were incubated with FITC-HSA (Abcam, ab8030) at 37 C° for 2 h before SEC separation.

### Nanoparticle tracking analysis

Particle size and concentration were measured via nanoparticle tracking analysis (NTA) using NanoSight NS500 equipped with NTA 3.2 analytical software (Malvern Panalytic, UK). Briefly, samples were diluted in 200 nm-filtered PBS if required and acquired using the following settings: five 30-s videos were recorded per sample with a camera level of 13. Software settings for analysis were kept constant for every measurement (screen gain 20, minimum track length 3).

### Western blotting

EVs (2×10^9^ in 24 µL) were mixed with 8 µL of sample buffer (0.5 M dithiothreitol, 0.4 M sodium carbonate, 8% sodium dodecyl sulfate and 10% glycerol) and heated at 70 °C for 10 min. The mixture was loaded onto a NuPAGE Novex 4–12% Bis-Tris Protein Gel (Invitrogen, NP0335BOX) and separated at 120 V in NuPAGE MES SDS running buffer (Invitrogen, NP0002) for 2 h. Proteins on the gel were transferred to an iBlot nitrocellulose membrane (Invitrogen, IB23001) for 7 min using the iBlot system. Membranes were blocked with Odyssey blocking buffer (LI-COR, 927-60004) for 1 h under gentle shaking. Afterwards, the membrane was incubated overnight at 4°C with primary antibody solution (1:1000 dilution for anti-TSG101 [Abcam, ab30871], anti-Calnexin [ThermoFisher, PA5-19169] and anti-Syntenin-1 [Origene, TA504796]; 1:200 dilution for anti-CD81 [SantaCruz, sc-9158] and 1:10,000 dilution for anti-β-actin [Sigma, A5441]). The membrane was rinsed with PBS supplemented with 0.1% Tween 20 (PBS-T) for 3 times over 15 min and incubated with the corresponding secondary antibody (1:15,000 dilution for all, LI-COR) for 1 h. Membranes were rinsed with PBS-T for 3 times over 15 min, one time with PBS and visualized on the Odyssey infrared imaging system (LI-COR, US).

### Proteomics analysis

EVs produced by wildtype (WT) or transduced stable cell lines were subjected to proteomic analysis. Briefly, samples were run with ThermoFisher Scientific Q Exactive Plus LC-MS/MS with 1 hour gradient and analyzed using R (version 4.1.2, 2021-11-01), RStudio (version 2022.07.1+554) and DEP (version 1.16.0). After filtering out bovine serum-derived proteins, keratin, mNG and puromycin resistance proteins, a total of 2340 proteins across all samples were included for downstream analysis.

### Cryo-electron microscopy

4 µl of sample was adsorbed onto holey carbon-coated grid (Quantifoil, Germany) that was glow-discharged. After blotting with filter paper, the grid was vitrified into liquid ethane at −178 °C using a Vitrobot (FEI, Netherlands). The frozen grid was then transferred onto a Philips CM200-FEG electron microscope (FEI, Netherlands) using a Gatan 626 cryo-holder (GATAN Inc, USA). Electron images were acquired using a low-dose system (accelerating voltage of 200 kV; nominal magnification of 50,000; temperature of −175°C). Defocus values ranged from −2 µm to −3 µm. Micrographs were recorded using a CMOS camera (TVIPS, Germany) at 4K × 4K.

### Confocal microscopy

Huh-7 cells were seeded at 10,000 per well in glass-bottom chamber slides (ThermoFisher; 155409) and incubated overnight. The cells were treated with HiBiT-mNG-labeled EVs (1×10^7^/µL) and incubated for 3.5 h. Hoechst 33342 (ThermoFisher, H3570) and LysoTracker Red DND-99 (ThermoFisher, L7528) was added to visualize nuclei and lysosomes, respectively. At 4 h, chamber slides were transferred to a microscope stage-top incubator at 37 °C with 5% CO_2_. Imaging was carried out using a confocal microscope (A1R confocal, Nikon, Japan) and analyzed by the NIS-Elements software (Nikon, Japan). 3D reconstruction was compiled with a height of 10 μm and a resolution of 0.3 μm.

### Flow cytometry for cells

Huh-7 cells were seeded at 3 ×10^4^ cells per well in 96-well plate and incubated overnight. The cells were treated with mNG-HiBiT-labeled EVs for 8 h. HUVECs were seeded at 5 ×10^4^ cells per well in 24-well plates and incubated overnight. The cells were stimulated with TNF-a (20 ng/mL) for 2 h and treated with mNG-HiBiT-labeled EVs for another 6 h. After trypsinization, cells were resuspended in 100 µL of PBS containing 2% FBS. 4ʹ,6-diamidino-2-phenylindole (DAPI) was added to all samples to exclude dead cells. The samples were measured with MACSQuant Analyzer 10 cytometer (Miltenyi, Germany). Data was analyzed with FlowJo software (version 10.6.2) and doublets were excluded by forward scatter area versus height gating.

### Flow cytometry for multiplex beads

MACSPlex Exosome Kit (Miltenyi Biotec; 130-108-813) was used to characterize the surface protein composition of EVs following manufacturers’ instructions. In brief, EVs (1 × 10^9^ in 120 µL) were incubated with 15 µL of MACSPlex exosome capture beads overnight in wells of a pre-wet and drained MACSPlex 96-well 0.22 µm filter plate at room temperature. The beads were rinsed with 200 µL MACSPlex buffer and detected after staining with APC-conjugated antibody mixture (anti-CD9/CD63/CD81) or AF647-conjugated anti-TSPAN2 antibodies (FAB7876R; R&D systems) for 1 h at room temperature. Next, the samples were rinsed twice, resuspended and analyzed using MACSQuant Analyzer 10 flow cytometer. FlowJo (v.10.6.2) was used to analyze data. Median fluorescence intensities (MFIs) for all 39 capture bead subsets were background-corrected by subtracting the respective values from matched non-EV-containing buffer controls and normalized to the beads with the highest level.

### Flow cytometry for single vesicle

mNG-HiBiT-labeled EVs were analyzed at a single-vesicle level on an Amnis CellStream instrument (Luminex) equipped with 405, 488, 561 and 642 nm lasers based on previously optimized settings and protocols^39^ and as described recently^58^ In brief, EV samples at a concentration of 1 × 10^10^ particles/mL were incubated with AF647-labelled anti-TSPAN2 antibody (R&D systems, FAB7876R-100UG, 0.5 µL) or a mixture of APC-labelled anti-CD9 (Miltenyi Biotech, clone SN4), anti-CD63 (Miltenyi Biotec, clone H5C6) and anti-CD81 antibodies (Beckman Coulter, clone JS64) at a concentration of 8 nM overnight and diluted by 2,000-fold in PBS-HAT buffer before data acquisition. Samples were measured with FSC turned off, SSC laser set to 40%, and all other lasers set to 100%. EVs were defined as SSC (low) by using mNG-tagged EVs as biological reference material, and regions to quantify mNG^+^ or AF647/APC^+^ fluorescent events were set according to unstained non-fluorescent samples and single fluorescence positive mNG-tagged reference EV controls. Data was analyzed using FlowJo software (version 10.6.2).

### Animal experiments

All animal procedures were performed in accordance with the ethical permission granted by Swedish Jordbruksverket. To check biodistribution of Tluc-labeled EVs, female NMRI mice with a bodyweight of around 20 g were administered intraperitoneally with 150 mg/kg D-luciferin (PerkinElmer, 122799). After 5 min, EVs (containing the same amount of Tluc in 100 µL) were injected through the tail vein. Live animals (isoflurane sedated) were imaged every 5 min over 30 min by IVIS Spectrum (PerkinElmer, US) with an exposure time of 30 seconds. Immediately after completion of IVIS session, the mouse was bled for collecting blood in EDTA-coated tubes and sacrificed for collecting major organs. Blood samples were immediately centrifuged at 2000 g for 10 min to retrieve plasma. The organs were weighed and lysed in 1 mL Triton X-100 solution (0.1% in PBS) using a TissueLyser II (QIAGEN, Germany) according to the manufacturer protocol. To analyze plasma retention of EVs, Nluc-labeled EVs were injected through tail vein. Blood samples were taken 60 min and 270 min after injection to retrieve plasma.

### Luciferase detection assay

Tluc was quantified in five types of samples: cell lysate, conditioned media, SEC eluate, mouse plasma and tissue lysate. To liberate Tluc from cells, the cell pellet was suspended in 100 µL Triton X-100 solution (0.1% in PBS) and shaken horizontally at 500 RPM for 10 min. Mouse plasma and tissue lysate were diluted by 5-fold and 10-fold with Triton X-100 solution, respectively. Typically, 25 μL of samples was added into white-walled 96-well plates and equal volume of ready-to-use Tluc substrate (Promega; E1501) was injected to each well. The luciferase intensity in each well was immediately measured using a GloMax 96 Microplate Luminometer machine (Promega, US). For conditioned media, 25 μL of samples was mixed with 25 μL PBS or Triton X-100 solution. The plate was shaken horizontally at 500 RPM for 5 min before addition of Tluc substrate. To discern resistant protein aggregate in SEC eluate, 25 µL of samples was mixed with Triton X-100 solution and then incubated with 25 µL of Proteinase K (ProK; Qiagen, 19131; 100 µg/mL in PBS) at 37°C for 30 min prior to Tluc measurement.

To detect Nluc in conditioned media and plasma, 25 μL of samples was added into white-walled 96-well plates along with 25 µL of Triton X-100 solution. The plate was shaken horizontally at 500 RPM for 5 min before addition of 25 µL of ready-to-use Nano-Glo substrate (Promega; N1130) for measuring luciferase intensity.

To detect HiBiT in conditioned media, 25 μL of samples was added into white-walled 96-well plates along with 25 µL PBS or Triton X-100 solution. The plate was shaken horizontally at 500 RPM for 5 min. 50 µL of ready-to-use HiBiT Lytic Detection mixture (Promega; N3040) was added to each well. After incubation at room temperature under horizontal shaking at 500 RPM for 10 min, the plate was immediately measured.

### Statistical analysis

All data were shown as mean ± standard deviation if applicable. GraphPad Prism (version 9.2.0) was used for statistical analysis and graph plotting.

## ACKNOWLEDGMENT

We thank Dave Carter, David Lowe, Tony De Fougerolles for editing the manuscript. S.E.-A. is supported by H2020 EXPERT, the Swedish foundation of Strategic Research (SSF-IRC; FormulaEx), ERC CoG (DELIVER) and the Swedish Medical Research Council. A.G. is an International Society for Advancement of Cytometry (ISAC) Marylou Ingram Scholar 2019-2023.

## AUTHOR CONTRIBUTION

DG, JZN and Spara EA devised the study. WZ, JR, DG, JZN and SEA designed the experiments and were involved in the discussion of results. WZ, JR, AG, SR and JB performed the *in vitro* experiments. WZ, JR and YZ performed the *in vivo* experiments. All authors were involved in writing and editing the manuscript.

## REFERENCE

1. Van Niel, G., D’Angelo, G. & Raposo, G. Shedding light on the cell biology of extracellular vesicles. Nature Reviews Molecular Cell Biology vol. 19 213–228 Preprint at https://doi.org/10.1038/nrm.2017.125 (2018).

2. Wiklander, O. P. B., Brennan, M., Lötvall, J., Breakefield, X. O. & Andaloussi, S. E. L. Advances in therapeutic applications of extracellular vesicles. Sci Transl Med 11, 1–16 (2019).

3. Couch, Y. et al. A brief history of nearly EV-erything - The rise and rise of extracellular vesicles. J Extracell Vesicles 10, (2021).

4. Herrmann, I. K., Wood, M. J. A. & Fuhrmann, G. Extracellular vesicles as a next-generation drug delivery platform. Nat Nanotechnol 16, 748–759 (2021).

5. Cheng, L. & Hill, A. F. Therapeutically harnessing extracellular vesicles. Nat Rev Drug Discov 0123456789, (2022).

6. Lindenbergh, M. F. S. & Stoorvogel, W. Antigen Presentation by Extracellular Vesicles from Professional Antigen-Presenting Cells. Annu Rev Immunol 36, 435–459 (2018).

7. Fu, W. et al. CAR exosomes derived from effector CAR-T cells have potent antitumour effects and low toxicity. Nat Commun 10, (2019).

8. Wu, C. H. et al. Extracellular vesicles derived from natural killer cells use multiple cytotoxic proteins and killing mechanisms to target cancer cells. J Extracell Vesicles 8, (2019).

9. Pelissier Vatter, F. A., et al. Extracellular vesicle- And particle-mediated communication shapes innate and adaptive immune responses. Journal of Experimental Medicine 218, 1–14 (2021).

10. Rani, S., Ryan, A. E., Griffin, M. D. & Ritter, T. Mesenchymal Stem Cell-derived Extracellular Vesicles: Toward Cell-free Therapeutic Applications. Molecular Therapy 23, 812 (2015).

11. Kooijmans, S. A. A. et al. Electroporation-induced siRNA precipitation obscures the efficiency of siRNA loading into extracellular vesicles. Journal of Controlled Release 172, 229–238 (2013).

12. Schulz-Siegmund, M. & Aigner, A. Nucleic acid delivery with extracellular vesicles. Adv Drug Deliv Rev 173, 89–111 (2021).

13. Liang, Y., Duan, L., Lu, J. & Xia, J. Engineering exosomes for targeted drug delivery. Theranostics 11, 3183–3195 (2021).

14. Kwon, S. et al. Engineering approaches for effective therapeutic applications based on extracellular vesicles. Journal of Controlled Release 330, 15–30 (2021).

15. Rädler, J., Gupta, D., Zickler, A. & Andaloussi, S. EL. Exploiting the biogenesis of extracellular vesicles for bioengineering and therapeutic cargo loading. Molecular Therapy 31, 1231–1250 (2023).

16. Richter, M., Vader, P. & Fuhrmann, G. Approaches to surface engineering of extracellular vesicles. Adv Drug Deliv Rev 173, 416–426 (2021).

17. Yim, N. et al. Exosome engineering for efficient intracellular delivery of soluble proteins using optically reversible protein-protein interaction module. Nat Commun 7, 1–9 (2016).

18. Gupta, D. et al. Quantification of extracellular vesicles in vitro and in vivo using sensitive bioluminescence imaging. J Extracell Vesicles 9, 1800222 (2020).

19. Joshi, B. S., de Beer, M. A., Giepmans, B. N. G. & Zuhorn, I. S. Endocytosis of Extracellular Vesicles and Release of Their Cargo from Endosomes. ACS Nano 14, 4444–4455 (2020).

20. Choi, H. et al. Exosome-based delivery of super-repressor IκBα relieves sepsis-associated organ damage and mortality. Sci Adv 6, 1–10 (2020).

21. Gupta, D., et al. Amelioration of systemic inflammation via the display of two different decoy protein receptors on extracellular vesicles. Nat Biomed Eng 5, (2021).

22. Liang, X. et al. Multimodal engineering of extracellular vesicles for efficient intracellular protein delivery. bioRxiv 2023.04.30.535834 (2023) doi:10.1101/2023.04.30.535834.

23. Kojima, R. et al. Designer exosomes produced by implanted cells intracerebrally deliver therapeutic cargo for Parkinson’s disease treatment. Nat Commun 9, (2018).

24. Gee, P. et al. Extracellular nanovesicles for packaging of CRISPR-Cas9 protein and sgRNA to induce therapeutic exon skipping. Nat Commun 11, 1334 (2020).

25. Fabbiano, F. et al. RNA packaging into extracellular vesicles: An orchestra of RNA-binding proteins? J Extracell Vesicles 10, (2020).

26. Zickler, A. M. et al. Novel endogenous engineering platform for robust loading and delivery of functional mRNA by extracellular vesicles. bioRxiv 2023.03.17.533081 (2023) doi:10.1101/2023.03.17.533081.

27. Willms, E., Cabañas, C., Mäger, I., Wood, M. J. A. & Vader, P. Extracellular vesicle heterogeneity: Subpopulations, isolation techniques, and diverse functions in cancer progression. Front Immunol 9, 738 (2018).

28. Kalluri, R. & LeBleu, V. S. The biology, function, and biomedical applications of exosomes. Science 367, (2020).

29. Hurwitz, S. N. et al. Proteomic profiling of NCI-60 extracellular vesicles uncovers common protein cargo and cancer type-specific biomarkers. Oncotarget 7, 86999–87015 (2016).

30. Jankovičová, J., Sečová, P., Michalková, K. & Antalíková, J. Tetraspanins, More than Markers of Extracellular Vesicles in Reproduction. Int J Mol Sci 21, 7568 (2020).

31. Corso, G. et al. Systematic characterization of extracellular vesicles sorting domains and quantification at the single molecule–single vesicle level by fluorescence correlation spectroscopy and single particle imaging. J Extracell Vesicles 8, 1663043 (2019).

32. Dooley, K. et al. A versatile platform for generating engineered extracellular vesicles with defined therapeutic properties. Molecular Therapy 29, 1729–1743 (2021).

33. Silva, A. M. et al. Quantification of protein cargo loading into engineered extracellular vesicles at single-vesicle and single-molecule resolution. J Extracell Vesicles 10, (2021).

34. Haraszti, R. A. et al. High-resolution proteomic and lipidomic analysis of exosomes and microvesicles from different cell sources. J Extracell Vesicles 5, (2016).

35. Böing, A. N. et al. Single-step isolation of extracellular vesicles by size-exclusion chromatography. J Extracell Vesicles 3, (2014).

36. Takakura, Y., Matsumoto, A. & Takahashi, Y. Therapeutic application of small extracellular vesicles (SEVs): Pharmaceutical and pharmacokinetic challenges. Biol Pharm Bull 43, 576–583 (2020).

37. Gao, Y. et al. Small Extracellular Vesicles: A Novel Avenue for Cancer Management. Front Oncol 11, 1–20 (2021).

38. Bost, J. P. et al. Growth Media Conditions Influence the Secretion Route and Release Levels of Engineered Extracellular Vesicles. Adv Healthc Mater 11, 1–15 (2022).

39. Görgens, A. et al. Optimisation of imaging flow cytometry for the analysis of single extracellular vesicles by using fluorescence-tagged vesicles as biological reference material. J Extracell Vesicles 8, 1587567 (2019).

40. Tertel, T. et al. High-Resolution Imaging Flow Cytometry Reveals Impact of Incubation Temperature on Labeling of Extracellular Vesicles with Antibodies. Cytometry A 97, 602–609 (2020).

41. Somiya, M. & Kuroda, S. Real-Time Luminescence Assay for Cytoplasmic Cargo Delivery of Extracellular Vesicles. Anal Chem 93, 5612–5620 (2021).

42. Trajkovic, K. et al. Ceramide triggers budding of exosome vesicles into multivesicular endosomes. Science 319, 1244–1247 (2008).

43. Wiklander, O. P. B. et al. Systematic methodological evaluation of a multiplex bead-based flow cytometry assay for detection of extracellular vesicle surface signatures. Front Immunol 9, 1326 (2018).

44. Liang, X. et al. Extracellular vesicles engineered to bind albumin demonstrate extended circulation time and lymph node accumulation in mouse models. J Extracell Vesicles 11, 11 (2022).

45. Zheng, W. et al. Cell-specific targeting of extracellular vesicles though engineering the glycocalyx. J Extracell Vesicles 11, (2022).

46. Yaseen, I. H., Monk, P. N. & Partridge, L. J. Tspan2: A tetraspanin protein involved in oligodendrogenesis and cancer metastasis. Biochem Soc Trans 45, 465–475 (2017).

47. Seipold, L. et al. Tetraspanin 3: A central endocytic membrane component regulating the expression of ADAM10, presenilin and the amyloid precursor protein. Biochim Biophys Acta Mol Cell Res 1864, 217–230 (2017).

48. Gonzales, P. A. et al. Large-Scale Proteomics and Phosphoproteomics of Urinary Exosomes. J Am Soc Nephrol 20, 363 (2009).

49. Bari, R. et al. Tetraspanins Regulate the Protrusive Activities of Cell Membrane. Biochem Biophys Res Commun 415, 619 (2011).

50. Umeda, R. et al. Structural insights into tetraspanin CD9 function. Nature Communications 2020 11:1 11, 1–11 (2020).

51. van Niel, G. et al. The Tetraspanin CD63 Regulates ESCRT-Independent and -Dependent Endosomal Sorting during Melanogenesis. Dev Cell 21, 708–721 (2011).

52. Edgar, J. R., Eden, E. R. & Futter, C. E. Hrs- and CD63-dependent competing mechanisms make different sized endosomal intraluminal vesicles. Traffic 15, 197–211 (2014).

53. Larios, J., Mercier, V., Roux, A. & Gruenberg, J. ALIX- and ESCRT-III–dependent sorting of tetraspanins to exosomes. J Cell Biol 219, (2020).

54. Mizenko, R. R. et al. Tetraspanins are unevenly distributed across single extracellular vesicles and bias sensitivity to multiplexed cancer biomarkers. J Nanobiotechnology 19, 1–17 (2021).

55. Zheng, W. et al. Cell-specific targeting of extracellular vesicles though engineering the glycocalyx. J Extracell Vesicles 11, 12290 (2022).

56. Liang, X. et al. Extracellular vesicles engineered to bind albumin demonstrate extended circulation time and lymph node accumulation in mouse models. J Extracell Vesicles 11, e12248 (2022).

57. Corso, G. et al. Reproducible and scalable purification of extracellular vesicles using combined bind-elute and size exclusion chromatography. Sci Rep 7, 1–10 (2017).

58. Görgens, A. et al. Identification of storage conditions stabilizing extracellular vesicles preparations. J Extracell Vesicles 11, e12238–e12238 (2022).

