## Supplementary Materials for "Identification of Novel Scaffold Proteins for Improved Endogenous Engineering of Extracellular Vesicles"

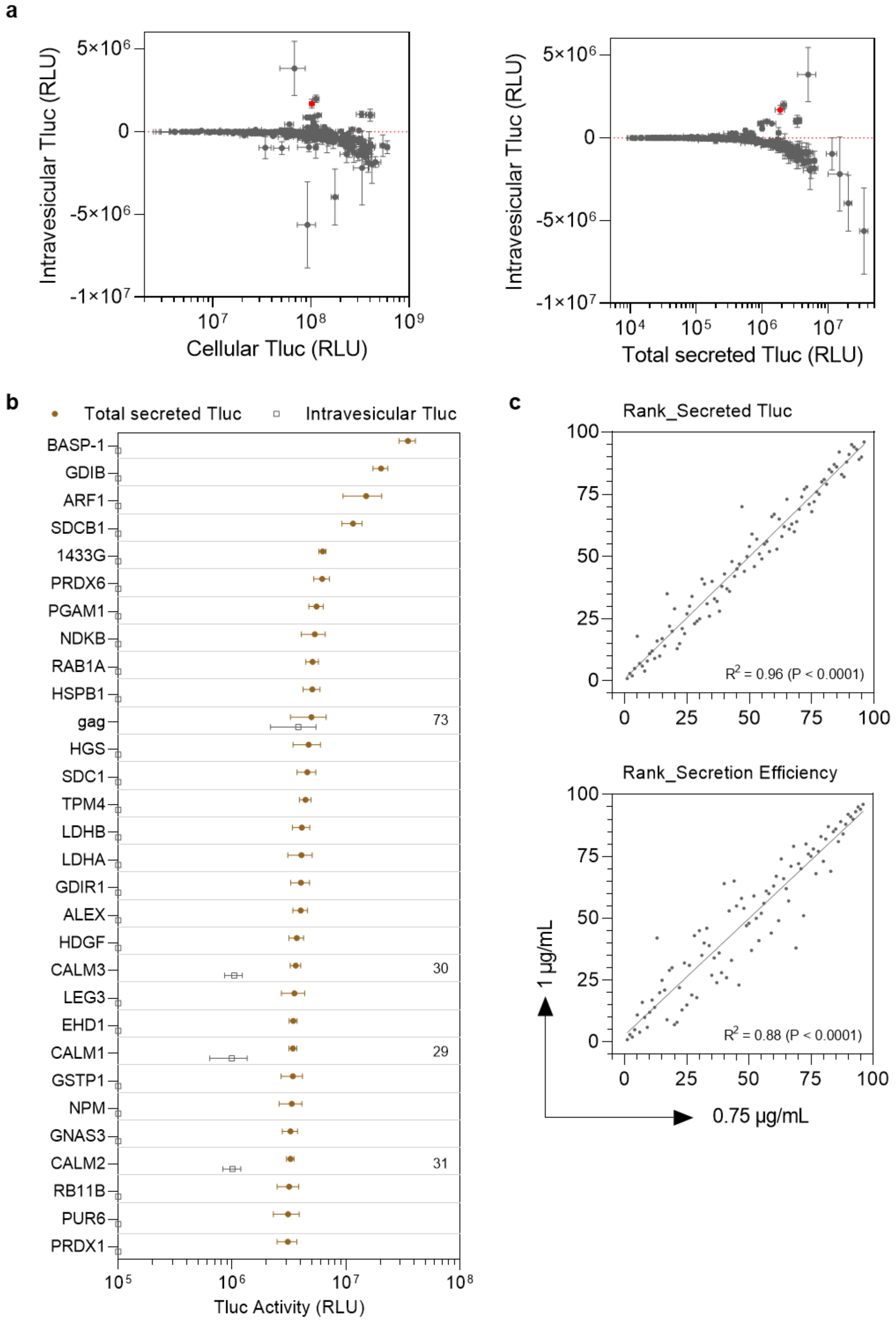

Figure S1. Screening results on HEK-293T cells. (a) Overview of intravesicular Tluc in relation to cellular Tluc and total secreted Tluc. (b) Top 30 proteins in terms of total secreted Tluc in the primary screening. Results were shown as the mean ± standard deviation of five biological replicates. Proteins are marked as gene name. The figure refers to the percent intravesicular Tluc. (b) Correlation regards the rank of secreted Tluc and secretion efficiency at different plasmid dose. Each dot refers to one candidate protein. Three biological replicates for each protein. The degree of correlation was performed using linear regression and shown as goodness-of-fit ($R^2$) and significance of none-zero slope (P).

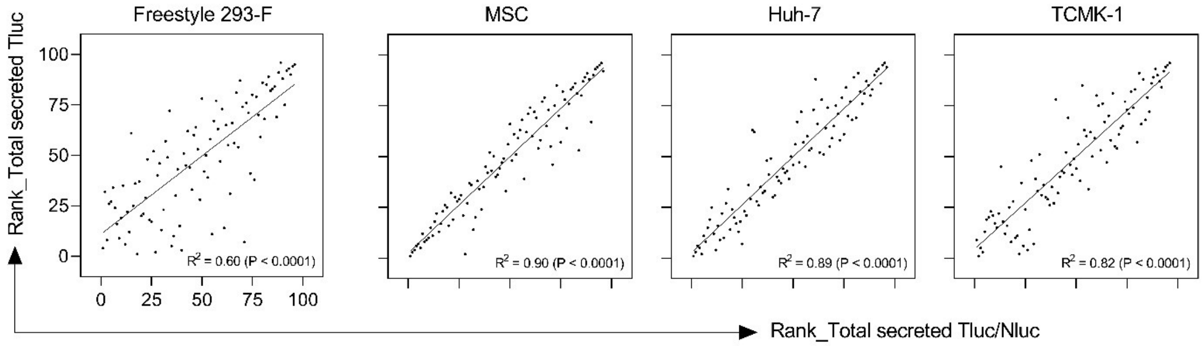


Figure S2. Correlation between the rank of total secreted Tluc and that of secreted Tluc/Nluc in different producer cell lines. Three biological replicates for each protein. The degree of correlation was performed using linear regression and shown as goodness-of-fit ($R^2$) and significance of none-zero slope (P).

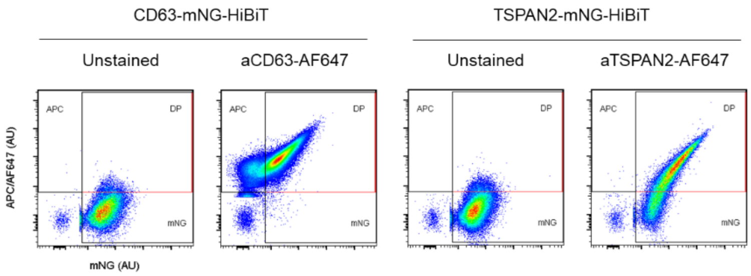


Figure S3. Detection of TSPAN2 and CD63 on engineered EVs using single-vesicle imaging flow cytometry.

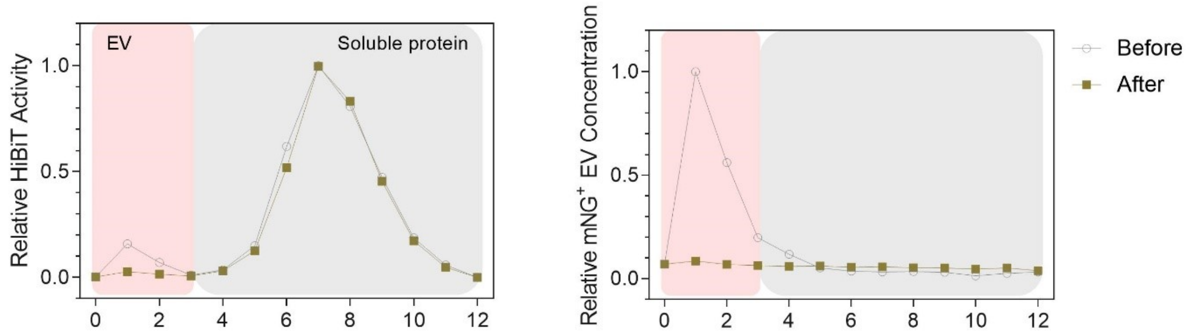


Figure S4. Effect of filtration on CALM1-engineered EVs. The conditioned media of CALM-HiBiT-mNG-transfected HEK-293T cells were directly fractionated or fractionated after 0.2 μm filtration. HiBiT activity and mNG$^+$ EV were quantified using bioluminescence and flow cytometry, respectively. Data were normalized to the fraction with maximum level.

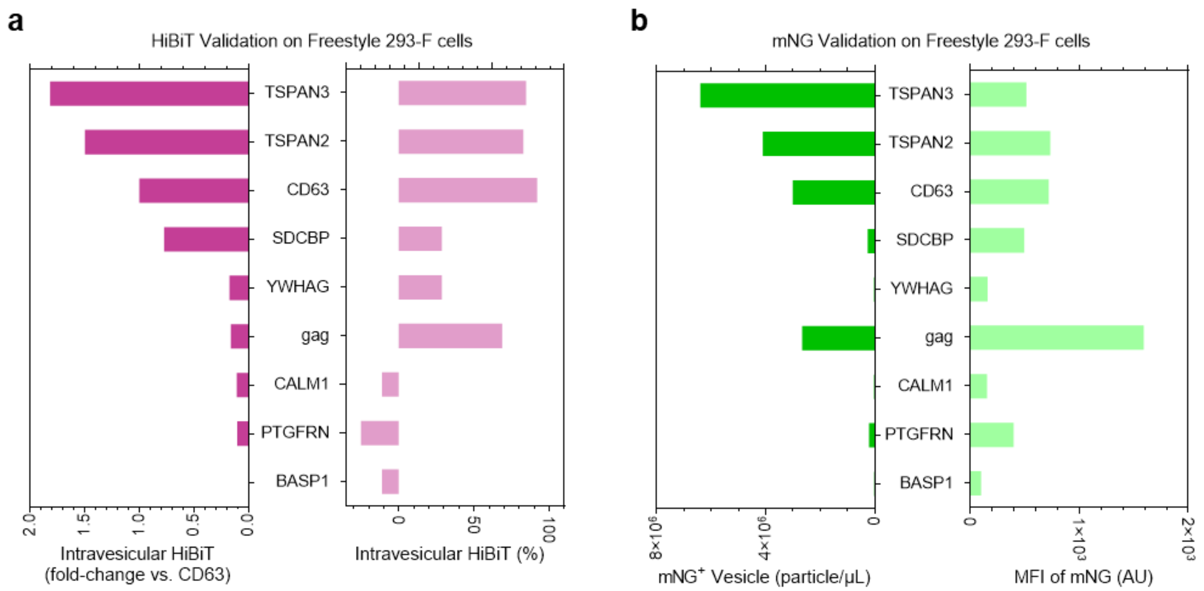


Figure S5. Quantification of HiBiT and mNG in the secretome of transfected Freestyle 293-F cells. Freestyle 293-F cells were grown in 6-well plate and transfected with 1.5 μg/mL plasmid for 48 hr. The conditioned media were pre-cleared and filtered through 0.2 μm membrane. (a) Quantification of intravesicular HiBiT and encapsulation index. (b) Quantification of the concentration and mean fluorescence intensity (MFI) of engineered EVs using single-vesicle flow cytometry. Proteins are marked as gene name.

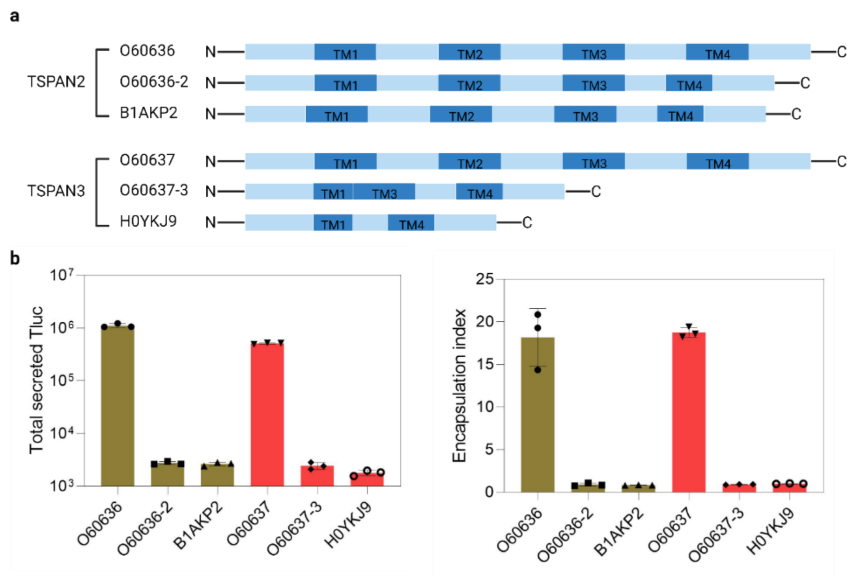


Figure S6. EV-sorting ability of TSPAN2 and TSPAN3 isoforms. (a) Topological scheme of isoforms. TM refers to transmembrane submain. Tluc was fused to the C-terminal of the isoforms. (b) Quantification of total secreted Tluc and encapsulation index in transfected HEK-293T cells. The cells were grown in 96-well microplate and transfected with 0.75 μg/mL plasmid for 48 hr. The cell cultures were centrifuged and Tluc activity was measured in the conditioned medium. Results were shown as the mean ± standard deviation of three biological replicates.

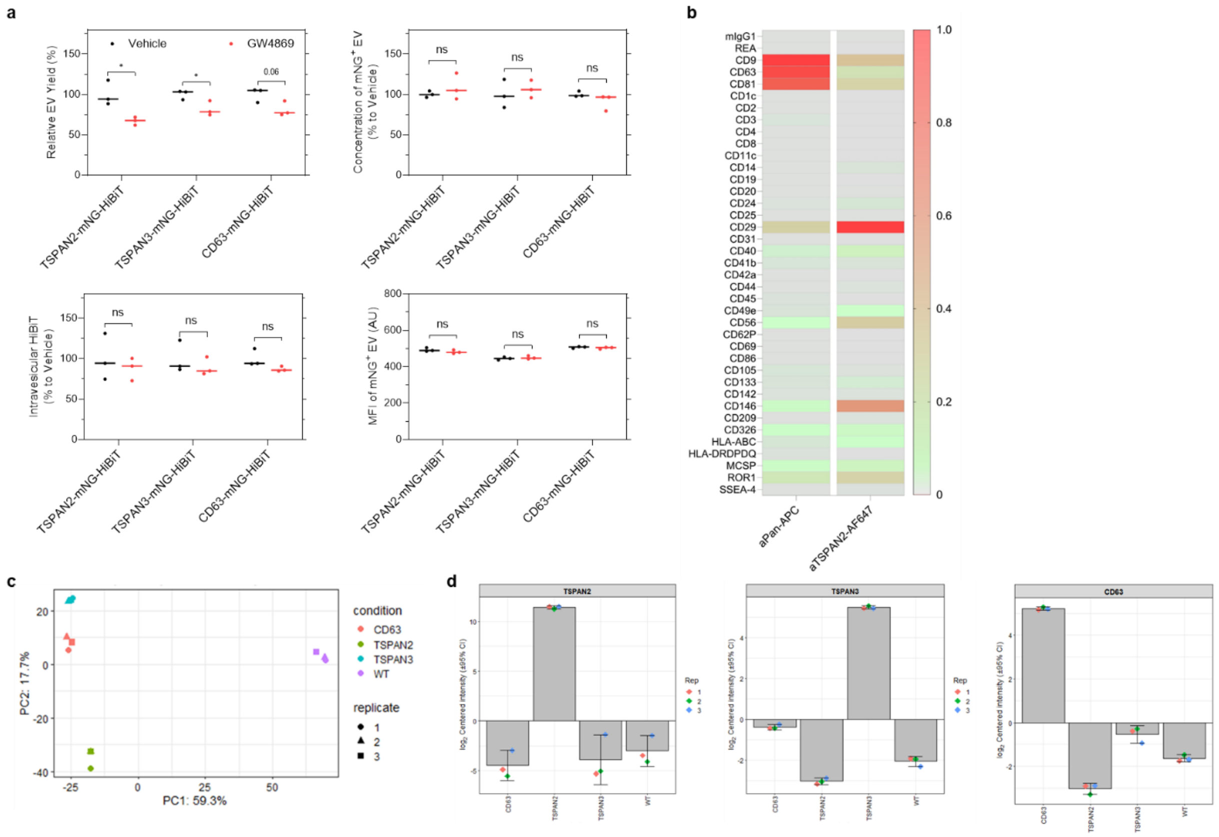


Figure S7. Physiochemical characterization of engineered EVs. (a) Effect of GW4869 on the yield of total and engineered EVs from transfected HEK-293T cells. GW4869 was added to Opti-MEM at a final concentration of 5 µm. (b) Surface epitope composition of EVs produced by TSPAN2-Tluc-transfected HEK -293T cells. EVs were captured with indicated beads (MACSPlex Exosome Kit, human), stained with Pan (CD9/CD63/CD81) or TSPAN2 detection antibodies, and detected by flow cytometry. (c) Principal clustering analysis of wildtype (WT) and engineered HEK-293T EVs using proteomics dataset. Engineered EVs were all labeled with mNG-HiBiT. (d) Relative level of TSPAN2, TSPAN3 and CD63 in EVs. EVs were collected from wildtype (WT) HEK-293T cells and cells stably expressing TSPAN-mNG-HiBiT. Quantification was based on the proteomics dataset in Figure 6f.

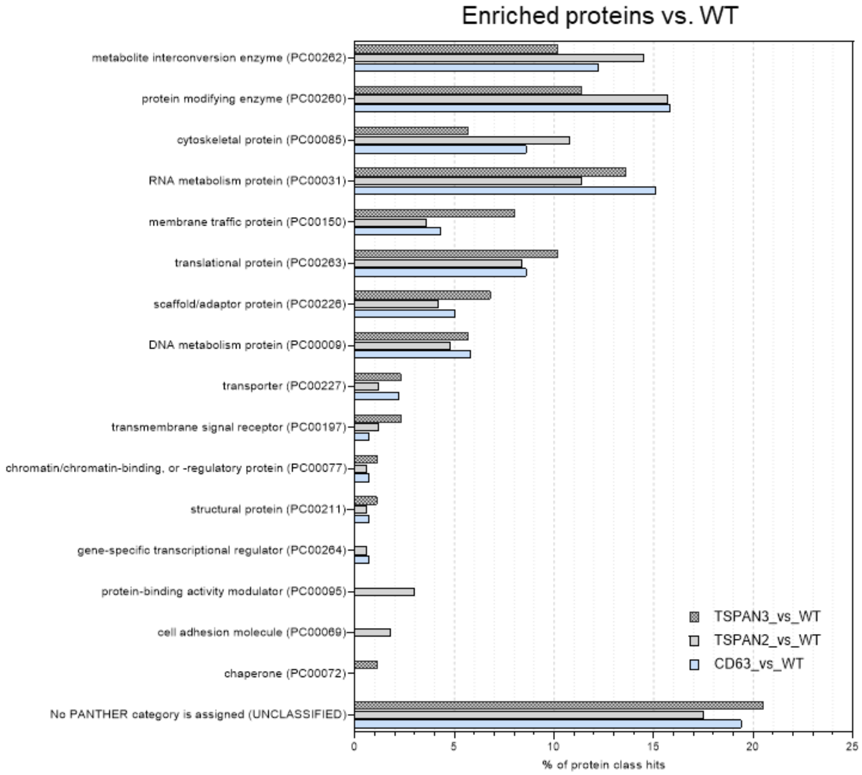


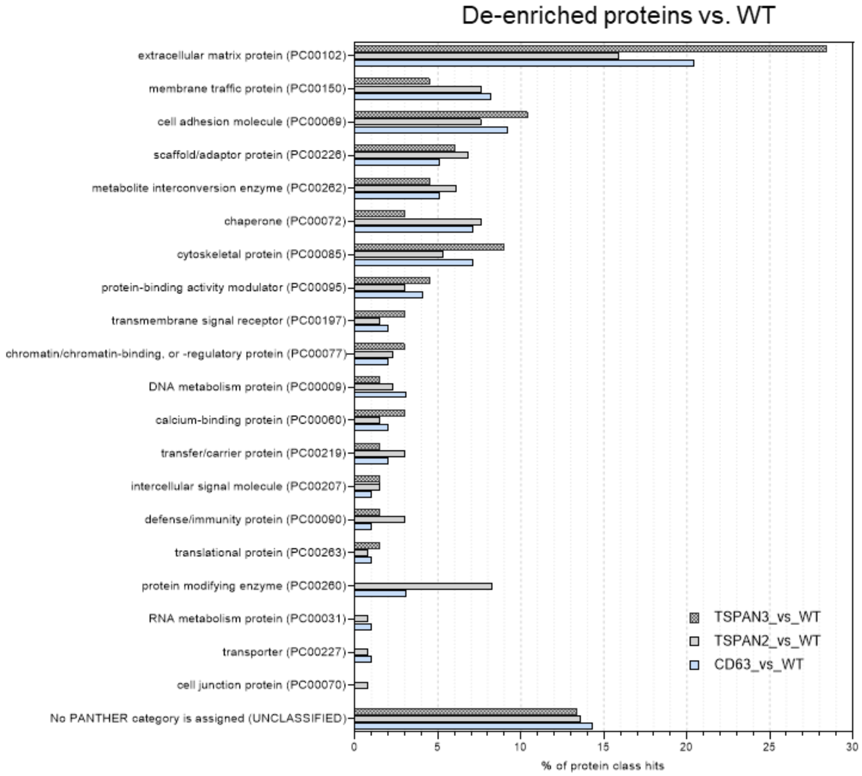


Figure S8. Gene ontology analysis of enriched and de-enriched proteins in engineered EVs compared to wildtype EVs.

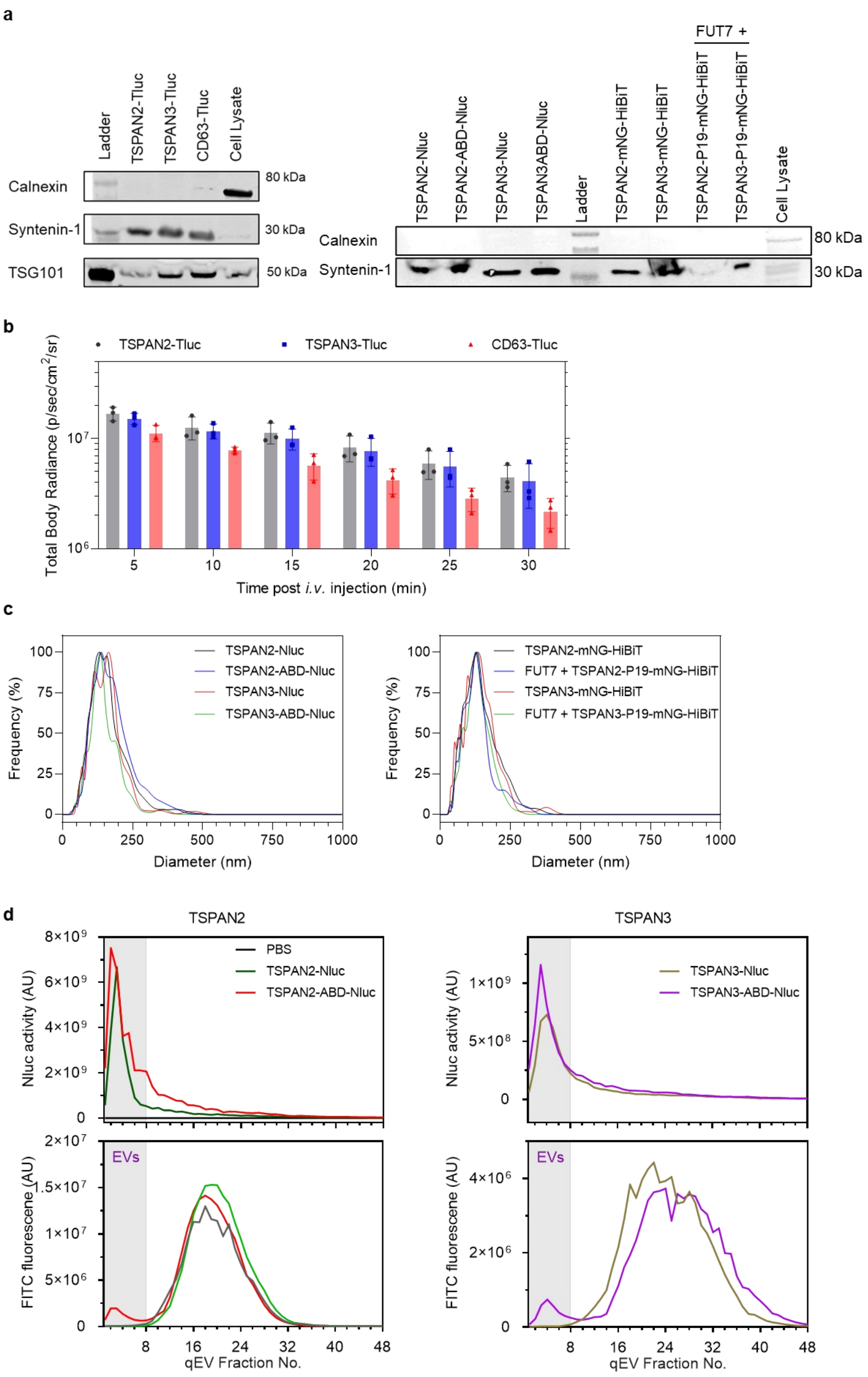
a
Ladder
TSPAN2-Tluc
TSPAN3-Tluc
CD63-Tluc
Cell Lysate
Calnexin
Syntenin-1
TSG101
80 kDa
30 kDa
50 kDa
FUT7 +
TSPAN2-Nluc
TSPAN2-ABD-Nluc
TSPAN3-Nluc
TSPAN3ABD-Nluc
Ladder
TSPAN2-mNG-HiBiT
TSPAN3-mNG-HiBiT
TSPAN2-P19-mNG-HiBiT
TSPAN3-P19-mNG-HiBiT
Cell Lysate
Calnexin
Syntenin-1
80 kDa
30 kDa
b
TSPAN2-Tluc
TSPAN3-Tluc
CD63-Tluc
Total Body Radiance (p/sec/cm2/sr)
Time post i.v. injection (min)
c
Frequency (%)
Diameter (nm)
TSPAN2-Nluc
TSPAN2-ABD-Nluc
TSPAN3-Nluc
TSPAN3-ABD-Nluc
TSPAN2-mNG-HiBiT
FUT7 + TSPAN2-P19-mNG-HiBiT
TSPAN3-mNG-HiBiT
FUT7 + TSPAN3-P19-mNG-HiBiT
d
TSPAN2
TSPAN3
PBS
TSPAN2-Nluc
TSPAN2-ABD-Nluc
TSPAN3-Nluc
TSPAN3-ABD-Nluc
Nluc activity (AU)
FITC fluorescene (AU)
EVs
qEV Fraction No.

Figure S9. Biological activity of engineered EVs. (a) Western blots of engineered EVs showing common EV markers and an exclusion marker. (b) Quantification of total body radiance of mice injected with Tluc labeled-EVs. Results are shown as mean ± standard deviation of three mice. (c) Size distribution of engineered EVs. Results are shown as the mean of five replicates. (d) Size exclusion chromatography elution profiles of Nluc-labeled EVs after incubating with FITC-HSA conjugates.
